# A graph-theoretical approach to DNA similarity analysis

**DOI:** 10.1101/2021.08.05.455342

**Authors:** Dong Quan Ngoc Nguyen, Lin Xing, Phuong Dong Tan Le, Lizhen Lin

## Abstract

One of the very active research areas in bioinformatics is DNA similarity analysis. There are several approaches using alignment-based or alignment-free methods to analyze similarities/dissimilarities between DNA sequences. In this work, we introduce a novel representation of DNA sequences, using *n*-ary Cartesian products of graphs for arbitrary positive integers *n*. Each of the component graphs in the representing Cartesian product of each DNA sequence contain combinatorial information of certain tuples of nucleotides appearing in the DNA sequence. We further introduce a metric space structure to the set of all Cartesian products of graphs that represent a given collection of DNA sequences in order to be able to compare different Cartesian products of graphs, which in turn signifies similarities/dissimilarities between DNA sequences. We test our proposed method on several datasets including Human Papillomavirus, Human rhinovirus, Influenza A virus, and Mammals. We compare our method to other methods in literature, which indicates that our analysis results are comparable in terms of time complexity and high accuracy, and in one dataset, our method performs the best in comparison with other methods.

## 1 Introduction

DNA similarity analysis is one of the main areas in bioinformatics. Two main approaches using in analyzing DNA sequences are *alignment-based methods* and *alignment-free methods*. Among the alignment-based methods, the multiple sequence alignment (MSA) method has the highest accuracy in analyzing similarities/dissimilarities between DNA sequences, but its time complexity increases extremely large for large datasets of DNA sequences whose lengths are sufficiently long. Thus searching for alignment-free methods that is effective in time complexity as well as having a reasonably high accuracy has been a very research problem in DNA similarity analysis. Alignment-free methods, using geometric approaches share a similar strategy that first embed DNA sequences into vectors in a Euclidean space, and then compute the similarity distance matrix, based on the underlying Euclidean distance, whose entries are distances between the representing vectors of DNA sequences. For papers containing alignment-free methods that use this approach, the reader is, for example, referred to [1, 2, 3, 4, 5, 6, 7, 8, 9, 10, 11, 12, 13, 14, 15, 16, 17, 18, 19, 20].

In this work, we propose an alignment-free method which avoids the embedding into Euclidean spaces and the use of numerical encoding of DNA sequences as vectors in a Euclidean space. The main observation in our approach is that every DNA sequence can be viewed as a string of letters–a combinatorial object in which each letter in the string is from the alphabet consisting of four nucleotides A, C, G, T. Using this observation, for an arbitrary positive integer *n*, and *n* positive integers *d*_1_, …, *d_n_*, our method allows to represent each DNA sequence as an *n*-ary Cartersian product of graphs, the *i*th component of which contains combinatorial information of *d_i_*-tuples of nucleotides appearing in the DNA sequence. In this way, the set of all *n*-ary Cartesian products of graphs can be viewed as an analogue of *n*-dimensional Euclidean spaces that contain the representing vectors of DNA sequences as in the traditional alignment-free methods. In order to be able to compare *n*-ary Cartesian products of graphs that represent DNA sequences in our proposed method, we use a variety of metric space structures on graphs that are available in geometric graph theory such as the metric space structures equipped with spectral distance metrics or matrix distance metrics. There are also other types of metric space structures on graphs (see [21]). Fixing, once and for all, a distance metric for each component in the set of all *n*-ary Cartesian products of graphs, we can equip this set with a metric space structure by simply taking the maximum of the values of all distances (see [22]). This procedure converts a given collection of DNA sequences into a metric space consisting of *n*-ary Cartesian products of graphs that contain combinatorial information of all *d_i_*-tuples of nucleotides in DNA sequences for any 1 ≤ *i* ≤ *n*, while also carrying a distance metric that allows to compare similarities/dissimilarities between *n*-ary Cartesian products of graphs that represent DNA sequences. The combinatorial information of tuples of nucleotides in DNA sequences can grow extremely large when allowing *n* to become sufficiently large. Thus in our approach, we choose suitable values of *n* to assure the fast time complexity while maintaining high accuracy in analyzing similarities/dissimilarities between DNA sequences. In comparison with other methods in literature such as the state-of-the-art Clustal Omega [23]–an MSA method, and the Fourier transform method in [24], an alignment-free method, our method perform comparable in accuracy and time complexity, and in some datasets (see Section 4), our method performs the best among all the methods with which we compare.

The structure of our paper is as follows. In Section 2, we introduce one-dimensional and high-dimensional graph representations of DNA sequences as well as their metric space structures that will be used in our experimental analysis. In Section 3, we describe our proposed method in detail. In Section 4, we apply our method to test on several real datasets including Human Papillomavirus (HPV) [25, 26], Human rhinovirus (HRV) [27], Influenza A virus [28, 29], and Mammals [30]. In the Appendix (see Section 5), we tabulate the GenBank ^*^ accession numbers of DNA sequences contained in the datasets on which we test our method.

## 2 Graph-theoretic representation of DNA sequences

In this section, we describe several representations of DNA sequences using graph theory, which allows to equip a given collection of graphs or Cartesian products of graphs with metric space structures. Such metric space structures of a collection of graphs will be exploited in our method for analyzing similarities/dissimilarities between DNA sequences.

### 2.1 One-dimensional Graph Representations

Let *α* denote a DNA sequence of length *m* of the form *a*_1_*a*_2_ · · · *a_m_*, where each *a_i_* is one of the nucleotides A, C, G, T. Let *d* be a positive integer such that *d* < *m*. In general, it suffices to choose small values of *d* between 2 and 10. Let *w* be a sufficiently small positive integer, and let *h* be the largest positive integer such that 0 ≤ *m* − (*wh* + 1) < *d* − 1. The last inequality condition assures that there are exactly *h d*-tuples of nucleotides appearing the sequence *α*, and the remaining nucleotides appearing after the *h*th *d*-tuple do not have enough *d* letters to form another *d*-tuple. Using such a pair of integers (*d, w*), we associate to *α* a *weighted undirected graph*, denoted by *G*_(*d,w*)_ whose nodes are constructed using *d* consecutive nucleotides in the sequence *α*, and two nodes form an edge when they are represented by two consecutive sequences of *d* nucleotides that are exactly *w* nucleotides apart from each other. For the rest of the paper, *w* is called the *window* of *α* that indicates the *distance* one needs to walk along the sequence *α* to construct nodes in *G*_(*d,w*)_. The number of nodes in *G*_(*d,w*)_ is at most *h*.

The first node, say *v*_1_, in *G*_(*d,w*)_ is represented by the ordered *d*-tuple *a*_1_*a*_2_ · · · *a_d_* consisting of the first *d* consecutive nucleotides in the DNA sequence *α*. In order to construct the second node, we start at nucleotide *a*_*w*+1_ which is exactly *w* nucleotides apart from *a*_1_, and form the second node *v*_2_ of the form *a*_*w*+1_*a*_*w*+2_ · · · *a*_*w*+*d*_. In general, by induction, we can define the *k*-th node, say *v_k_*, with 1 ≤ *k* ≤ *h*, as the ordered *d*-tuple *a*_*w*(*k*−1)+1_*a*_*w*(*k*−1)+2_ · · · *a*_*w*(*k*−1)+*d*_. Since each *a_i_* is one of the nucleotides A, C, G, T, there are only finitely many choices for each node *v_k_*. In fact, there are exactly 4*^d^* choices for each *v_i_*. So it may occur that the construction can result in *v_i_* = *v_j_* for some *i* ≢ *j*, which implies that the set of nodes of *G*_(*d,w*)_ consists of all the distinct elements in the multiset {*v*_1_, *v*_2_, …, *v_h_*}.

Two distinct nodes *v_i_, v_j_* form an edge *e* = (*v_i_, v_j_*) in *G*_(*d,w*)_ if there exists an integer 1 ≤ *k* ≤ *h* such that either of the following is true.

i. *v_i_* = *a_w_*(*k*−1)+1*a_w_*(*k*−1)+2 · · · *a_w_*(*k*−1)+*d* and *v_j_* = *a*_*wk*+1_*a*_*wk*+2_ · · · *a*_*wk*+*d*_.
ii. *v_j_* = *a_w_*(*k*−1)+1*a_w_*(*k*−1)+2 · · · *a_w_*(*k*−1)+*d* and *v_i_* = *a*_*wk*+1_*a*_*wk*+2_ · · · *a*_*wk*+*d*_.

In other words, *e* = (*v_i_, v_j_*) is represented by two consecutive ordered *d*-tuples appearing in the DNA sequence *α*. The *weight* of *e* is defined to be the number of times that the pair (*v_i_, v_j_*) that represents the edge *e*, appears as two consecutive ordered *d*-tuples in the sequence *α*.

#### Example 2.1

Let *α* denote the DNA sequence AACTGTATGACGTATG of length *m* = 16. We illustrate the above construction to represent *α* as a weighted undirected graph *G*_(2,1)_, where *d* = 2 and *w* = 1. Such a graph representation is called a *dinucleotide* representation with window 1. Using the above construction, we obtain that *v*_1_ = *AA*, *v*_2_ = *AC*, *v*_3_ = *CT*, *v*_4_ = *TG*, *v*_5_ = *GT*, *v*_6_ = *TA*, *v*_7_ = *AT*, *v*_8_ = *TG*, *v*_9_ = *GA*, *v*_10_ = *AC*, *v*_11_ = *CG*, *v*_12_ = *GT*, *v*_13_ = *TA*, *v*_14_ = *AT*, and *v*_15_ = *TG*. Thus the set of nodes of *G*_(2,1)_ consists of all distinct elements in the multiset {*v*_1_, *v*_2_, …, *v*_15_}, which implies that the set of nodes of *G*_(2,1)_ is AA, AC, AT, CG, CT, GA, GT, TA, TG. Note that *GT* and *TG* are distinct nodes since we consider the ordered 2-tuples appearing in *α*. The graph *G*_(2,1)_ that represents *α* is illustrated in Figure 1(A) in which the positive integer appearing on each edge indicates its weight. For example, the edge (AC, CG) appears in *G*_(2,1)_ with weight 1 since AC, CG appear as two consecutive ordered 2-tuples in *α* exactly one time.

**Figure 1:**
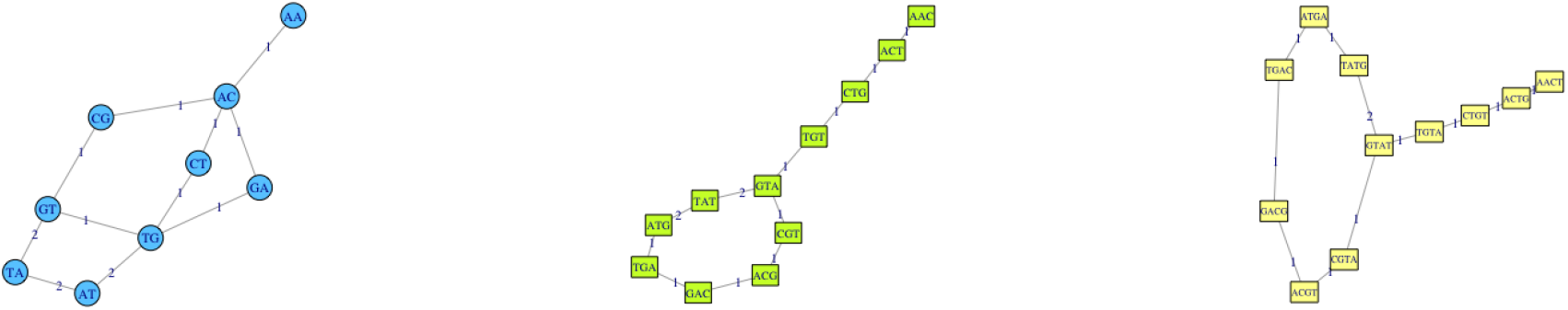
Graph representations of DNA sequence AACTGTATGACGTATG

When (*d, w*) is (3, 1) (resp., (4, 1), we, in a similar way as above, obtain the *trinucleotide* representation with window 1 (resp. the *tetranucleotide* representation with window 1) of *α*. See Figure 1(B) and (C) for the graphs *G*_(3,1)_ and *G*_(4,1)_.

### 2.2 High-dimensional Graph Representations

In this subsection, to each DNA sequence *α* we associate a Cartersian product of weighted undirected graphs. Let *α* be a DNA sequence of length *m*. Let (*d*_1_, *w*_1_), …,(*d_n_, w_n_*) be *n* pairs of positive integers, where each *d_i_* satisfies *d_i_* < *m*, and the *w_i_* are chosen to be sufficiently small. In our experimental analysis, we choose *w_i_* = 1, which is fast in time complexity as well as high in accuracy. For each (*d_i_, w_i_*) with 1 ≤ *i* ≤ *n*, we associate to *α* the weighted undirected graph 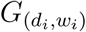 as in Subsection 2.1. We then combine all the graphs 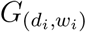, and associate to the DNA sequence *α* the *n*-ary Cartersian product 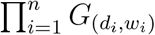, which can be viewed as a high-dimensional graph representation of *α*.

As illustration, let *α* be the DNA sequence as in Example 2.1. Let (*d*_1_*, w*_1_) = (2, 1), (*d*_2_*, w*_2_) = (3, 1), and (*d*_3_*, w*_3_) = (4, 1). Using the above construction, we obtain the 3-ary Cartersian product *G*_(2,1)_ × *G*_(3,1)_ × *G*_(4,1)_ that represents the sequence *α*, where *G*_(2,1)_, *G*_(3,1)_, and *G*_(4,1)_ are the graphs in Example 2.1 and Figure 1.

### 2.3 Finite Metric Structure of DNA Sequences

In this subsection, we introduce metric space structures to the Cartesian product representation of DNA sequences that is constructed in Subsection 2.2. Let 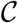 be a finite set of DNA sequences, the minimum of whose lengths is *m*. Let (*d*_1_*, w*_1_), …,(*d_n_, w_n_*) be *n* pairs of positive integers for some integer *n* ≥ 1, where each *d_i_* satisfies *d_i_* < *m*, and the *w_i_* are chosen to be sufficiently small. Let 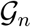 be the set of all *n*-ary Cartesian products of weighted undirected graphs. Using the construction in Subsection 2.2, for each DNA sequence 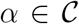, there is an *n*-ary Cartersian product 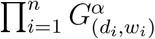 of weighted undirected graph that represents *α*. Thus we obtain a map 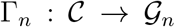 that sends each DNA sequence *α* to its associated *n*-ary Cartesian product 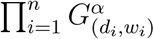. The map Γ*_n_* is called an *n-ary Cartersian product representation of* 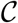.

It is well-known that there are many metric space structures for a collection of graphs (see, for example, [21]), i.e. for a given collection of graphs 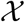, there exist several distance metrics 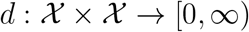 that satisfy the following conditions:

(D1) (**symmetry**) *d*(*G*_1_*, G*_2_) = *d*(*G*_2_*, G*_1_) for any graphs *G*_1_*, G*_2_ in 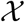; and
(D2) (**triangle inequality**) *d*(*G*_1_*, G*_2_) ≤ *d*(*G*_1_*, G*_3_) + *d*(*G*_2_*, G*_3_) for any graphs *G*_1_*, G*_2_*, G*_3_ in 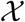.

In general, we do not require that *d*(*G*_1_*, G*_2_) = 0 if and only if *G*_1_ = *G*_2_, as in the traditional notion of a distance metric in mathematics (see, for example, [22]). In practice, and in our experimental analysis, we rarely encounter two graphs *G*_1_*, G*_2_ that represent two DNA sequences such that *d*(*G*_1_*, G*_2_) = 0. It suffices to obtain that *G*_1_ is very similar to *G*_2_ from the fact that *d*(*G*_1_*, G*_2_) is sufficiently small from which we wish to deduce that the DNA sequences that *G*_1_*, G*_2_ represent are similar.

For such a distance metric *d* satisfying (D1) and (D2) as above, we can equip the collection 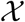 of graphs with a metric space structure associated to *d*, which allows us to compare similarities/dissimilarities between graphs contained in 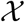. In this work, we use the distances for DNA similarity analysis that must scale linearly or near-linearly in the size in the graphs associated to the DNA sequences. We list below the distances that we will apply to our method. For the sake of brevity, we only cite the work that contains the detailed definitions of the distances as well as the reference containing the Python library that implements the graph distances used in this work.

1. **Adjacency spectral distance**: See [21, Section 2.2.1] for the description of this distance.
2. **Edit distance**: See [21, Equation (6), Section 2.2.2] for the description of this distance.

The Python library that implements the above distances used in this work can be accessed from the GitHub of Peter Wills ^*^.

Based on the well-known distance metrics on a collection of graphs, we can construct distance metrics for a collection of *n*-ary Cartersian products of graphs as follows. Let 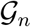 denote a collection of *n*-ary Cartersian products of graphs of the form 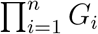 where the *G_i_* are weighted undirected graphs. For each 1 ≤ *i* ≤ *n*, let 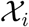 denote the collection of weighted undirected graphs *G_i_* appearing in the *i*-th component of the *n*-ary Cartersian products of graphs in 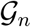. There is a natural bijection between 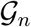 and the Cartesian product 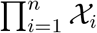.

For each 1 ≤ *i* ≤ *n*, take an arbitrary distance metric *d_i_* on 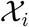; for example, one can choose *d_i_* to be either the adjacency spectral distance or the edit distance as above for each *i*. We can define a distance metric 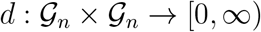 by setting

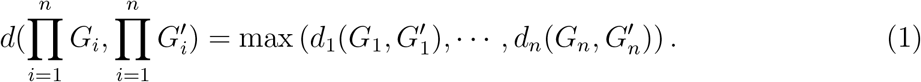

One can easily check that *d* satisfies the requirements of a distance metric, say (D1) and (D2) above. In our method, we will use the distance defined by (1) for DNA similarity analysis, using high-dimensional graph representation, where each *d_i_* is either the adjacency spectral distance or the edit distance.

## 3 Proposed Method

In this section, we describe our method for analyzing similarities/dissimilarities between DNA sequences. Our proposed method for reconstructing a phylogenetic tree of DNA sequences, using graph representations described in Section 2, is described in the following algorithm:

(0) (Input) A collection 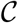 consisting of *m* DNA sequences *α*_1_*, …, α_m_*.
(1) Choose *n* tuples of positive integers (*d_i_, w_i_*) for 1 ≤ *i* ≤ *n*, where the *d_i_* is less than the minimum of the lengths of *α*_1_*, …, α_m_*, and the *w_i_* are sufficiently small. In practice, and in our experimental analysis, we choose *w_i_* = 1 for all *i*.
(2) Construct high-dimensional graph representation (HDGR) of each DNA sequence *α_i_* to obtain a finite collection 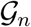 of *n*-ary Cartesian products of graphs 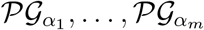, where the 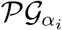 is the *n*-ary Cartesian product of graphs, corresponding to the DNA sequence *α_i_* (see Section 2.2).
(3) Associate the distance metric *d* on 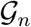 as in Subsection 2.3, where the *i*-th component distance metrics *d_i_* are chosen to be all adjacency spectral distances or edit distances. We use the Python library from the GitHub of Peter Wills ^†^ to implement the component distances, which will then be used to compute *d* by taking the maximum of the values of the component distances *d_i_*.
(4) Compute the distance matrix of dimensions *m × m* whose (*i, j*)-entry is the distance 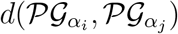.
(5) (Ouput) Construct the phylogenetic tree of the DNA sequences from the distance matrix in Step 3, using UPGMA algorithm (see [31]).

## 4 Experimental Analysis

In this section, we describe our experimental analysis, and the results we obtain from applying our proposed method in Section 3 to several real datasets including Human Papillomavirus (HPV) [25, 26], Human rhinovirus (HRV) [27], Influenza A virus [28, 29], and Mammals [30]. The GenBank ^‡^ accession numbers of DNA sequences contained in these datasets are listed in the Appendix (see Section 5). Computations in this research are implemented on a PC with configuration of Intel Core i7, CPU 2.50 GHz, and 16 GB 1600 MHz DDR3.

Each of the above datasets is represented by a finite collection 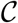 of DNA sequences *α*_1_, …, *α_l_* for various lengths *l*. We apply the proposed method in Section 3 for the collection 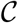. More precisely, in *Step 1*, we fix, once and for all, *n* = 3, (*d*_1_*, w*_1_) = (2, 1), (*d*_2_*, w*_2_) = (3, 1), and (*d*_3_*, w*_3_) = (4, 1). Thus in *Step 2*, we, for each *α_i_*, obtain a 3-ary Cartersian product of graphs 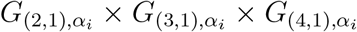 that represents *α_i_*. Here the construction of graphs 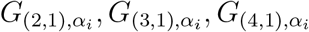 and their 3-ary Cartesian products follow Subsections 2.1 and 2.2. In other words, 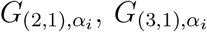, and 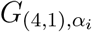 are dinucleotide, trinucleotide, and tetranucleotide representations with window 1 of *α_i_*, respectively. Thus we obtain a collection 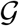 of exactly *l* 3-ary Cartesian products of graphs of the above form that represent all the *α_i_*. See Example 2.1 for an explicit example of a DNA sequence, and its graph representations.

In our computations, we try several representations of DNA sequences as in Section 2, but the results are appproximately similar, and the above representation provides us with the *fastest time complexity* and *highest accuracy* in analyzing similarities/dissimiarlities between DNA sequences. In *Step 3*, we use the distance metric *d* on 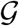 in Subsection 2.3, where the *i*-th component distance metrics *d_i_* are chosen to be all adjacency spectral distances or edit distances. Using the time complexity summary in [21], edit distances have faster time complexity than spectral distances. We also compare our results with other methods in literature such as the Fourier transform method developed in [24] and the state-of-the-art aligntment method for DNA similarity analysis called Clustal Omega in [23]. Our proposed method using edit distance performs the best in comparison with the two methods in [24] and [23] when applying to test on the above datasets. Tables 1 and 2 summarize time complexity and accuracy in applying our proposed methods, using both edit distances and adjacency spectral distances, to the datasets HPV, HRV, Influenza A virus, and Mammals. The tables also list time complexity and accuracy, using the methods in [24] and [23]. The computations and reconstruction of phylogenetic trees using Clustal Omega were produced in [24]. For the sake of brevity, we do not reproduce these computations in our paper, but instead refer the readers to [24] for the experimental analyses using the methods in [24] and [23].

**Table 1:** Time complexity using our proposed methods in comparison with those in [24] and [23].

| Method | Influenza | HPV | Mammals | HRV |
| --- | --- | --- | --- | --- |
| Edit distance | 0.51s (see Fig.2) | 43.31s (see Fig.4) | 4.34s (see Fig.6) | 8.15s (see Fig.8) |
| Spectral distance | 54.67s (see Fig.3) | 2h 29min (see Fig.5) | 42.60s (see Fig.7) | 10min (see Fig.9) |
| Clustal Omega in [23] | 9s | 2h 17min | 10min | 19min 35s |
| Fourier transform method in [24] | < 1s | < 30s | 4s | 7s |
| Number of DNA sequences | 38 | 400 | 31 | 116 |
| Lengths of DNA sequences | 1350 – 1467 | 7814 – 10424 | 16338 – 17447 | 6944 – 7458 |

**Table 2:** Accuracy using our proposed methods in comparison with those in [24] and [23].

| Method | Influenza | HPV | Mammals | HRV |
| --- | --- | --- | --- | --- |
| Edit distance | 1 (A/turkey/VA/H5N1)<sup>†</sup> (see Fig.2) | 1 (HPV11-14) (see Fig.4) | 0 (see Fig.6) | 0 (see Fig.8) |
| Spectral distance | 0 (see Fig.3) | 1 (HPV11-14) (see Fig.5) | 2 (Rabbit,Pig) (see Fig.7) | 1 (C c025*) (see Fig.9) |
| Clustal Omega in [23] | 1 (A/turkey/VA/H5N1)<sup>†</sup> | 0 | 1 (Squirrel) | 0 |
| Fourier transform method in [24] | 1 (A/duck/Guangxi/H1N1) | 1 (one from HPV11) | 1 (Squirrel) | 0 |

### 4.1 Influenza A virus

We first consider the dataset of Influenza A viruses. Influenza A viruses are very dangerous because they have a diverse range of hosts including birds, horses, swine, and humans. These viruses have been a serious health threat to humans and animals (see [32]), and are known to have high degree of genetic and antigenic variability (see [28, 29]). Some subtypes of Influenza A viruses are even lethal including H1N1, H2N2, H5N1, H7N3, and H7N9. We apply our method on the dataset consisting of 38 Influenza A virus genomes. From Figures 2, 3, and Tables 1 and 2, we find that our method, using spectral distance perform the best in terms of accuracry, in comparison with all other methods in the tables which incorrectly group one of the subtypes of Influenza A viruses. In term of time complexity, our method using edit distance is comparable with the Fourier transform method in [24], and both methods perform the best in terms of time complexity. Figures 2, 3 illustrate the phylogenetic trees of Influenza A viruses, based on our methods, using edit distance and spectral distance, respectively.

**Figure 2:**
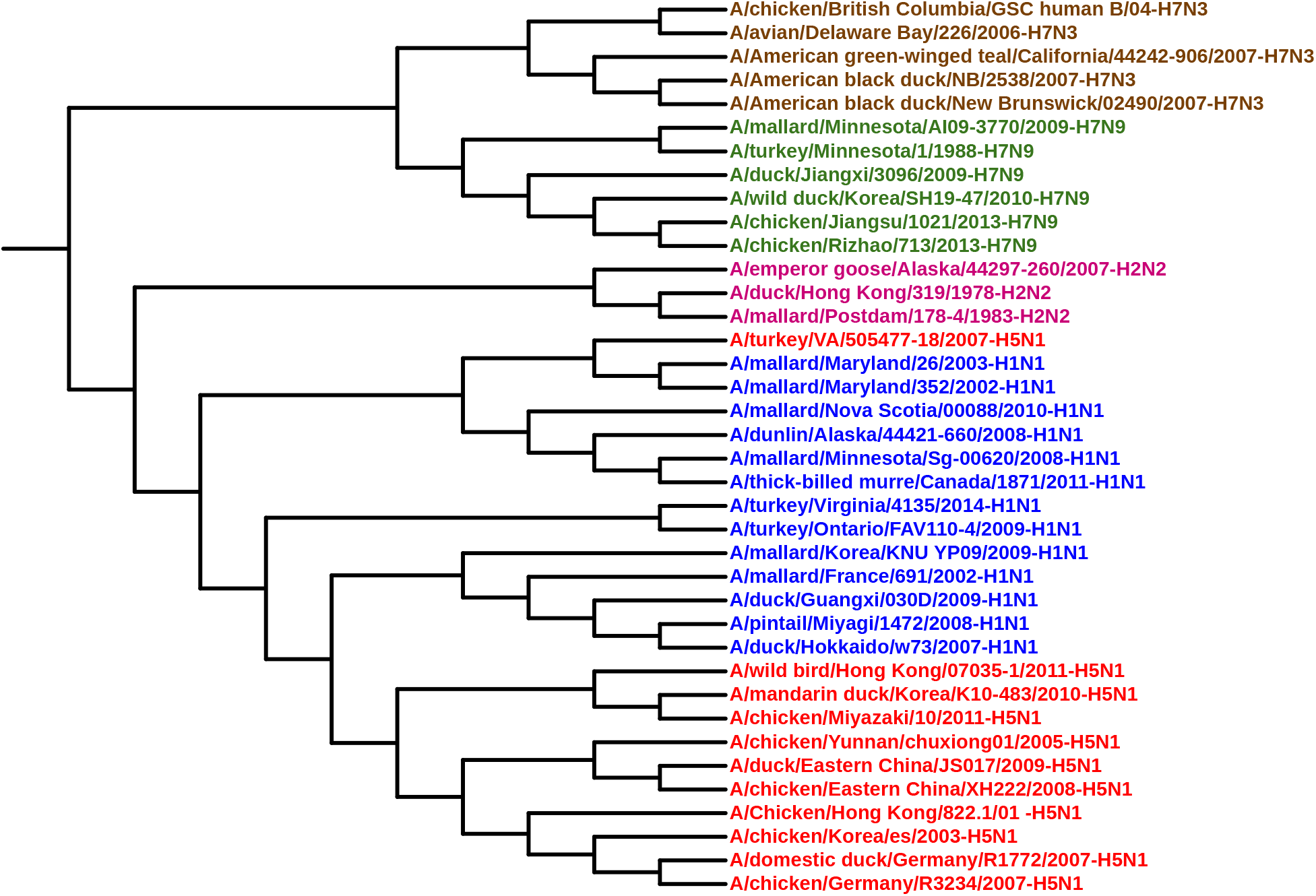
Phylogenetic tree of Influenza based on edit distance.

**Figure 3:**
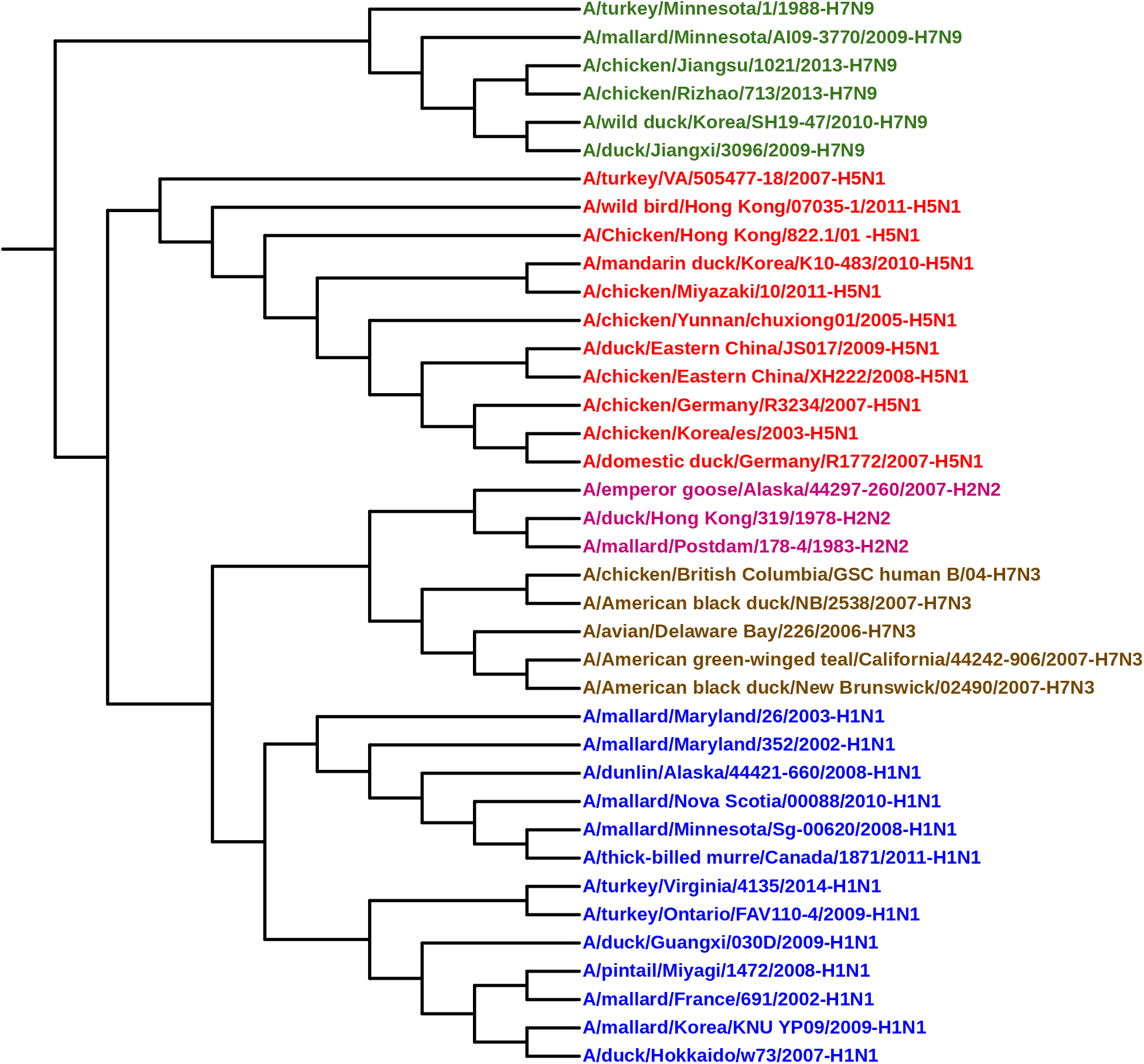
Phylogenetic tree of Influenza based on spectral distance.

### 4.2 Human Papillomavirus (HPV)

In this subsection, we consider the dataset of Human Papillomavirus (HPV). Human Papillomavirus is mostly responsible for cervical cancer which is the second most common cancer among women (see [25]). We apply our method, using both edit distance and adjacency spectral distance, on the data set of 400 HPV genomes. In terms of time complexity and accuracy, our method using edit distance is similar to the Fourier transform method in [24]. Both of our methods, using edit distance and spectral distance, incorrectly identify one HPV genome HPV11-14. In terms of time complexity, as mentioned above, edit distance is always faster than spectral distance. See Figures 4 and 5 for the phylogenetic trees of HPV, based on our methods, using edit distance and spectral distance, respectively.

**Figure 4:**
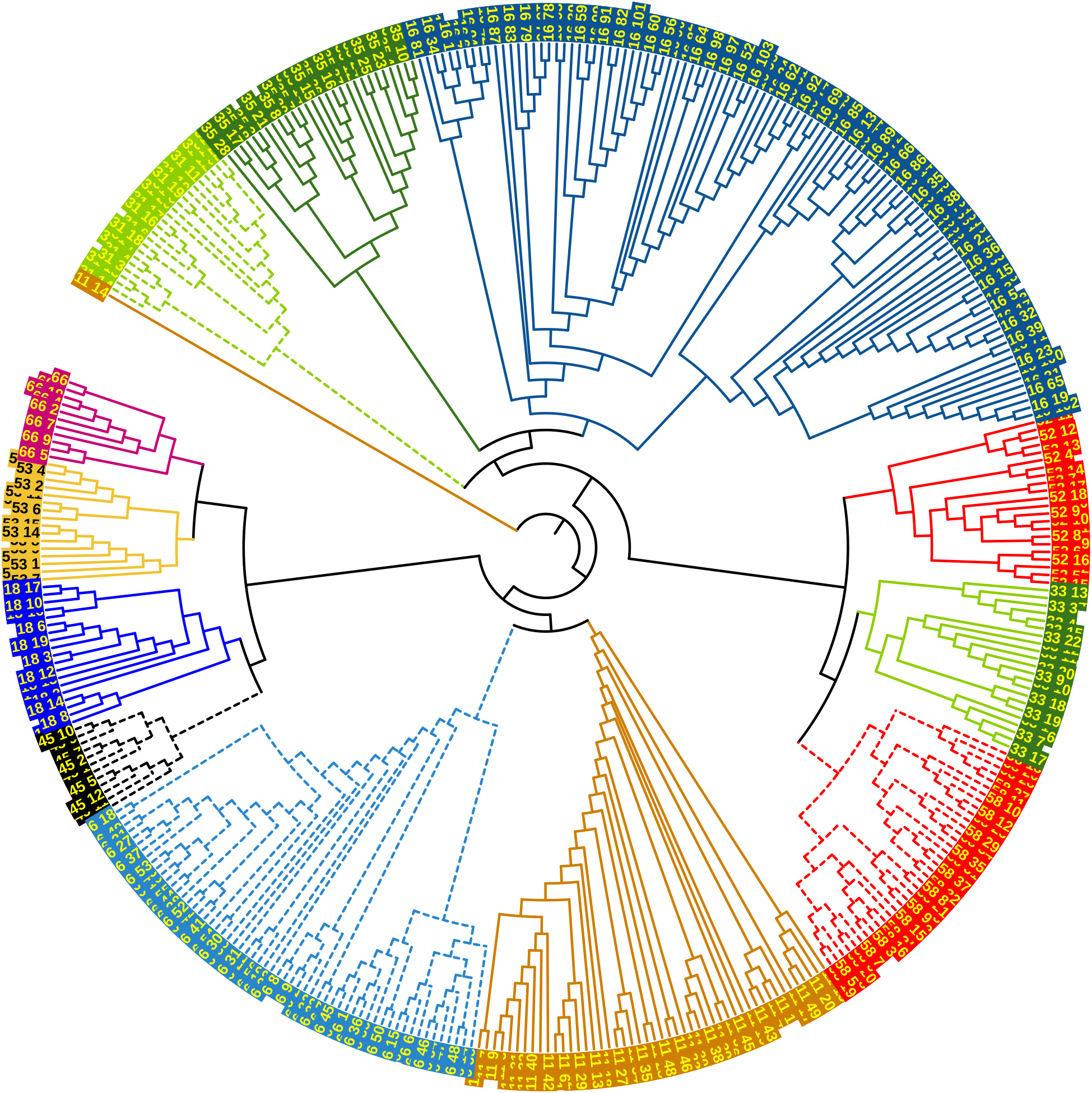
Phylogenetic tree of HPV based on edit distance.

**Figure 5:**
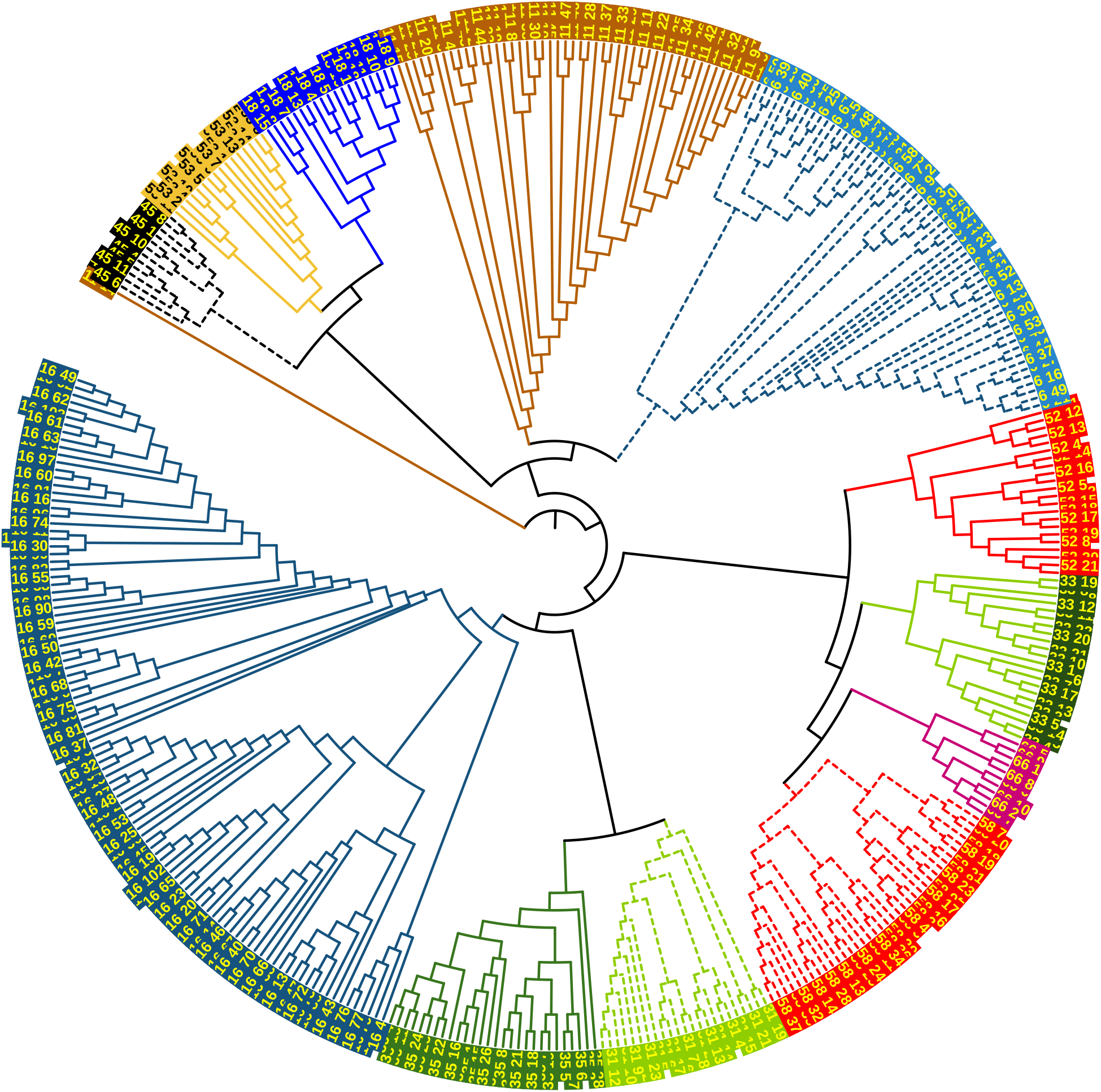
Phylogenetic tree of HPV based on spectral distance.

### 4.3 Mammals

It is known that there is a rapid mutation rate in the mitochondrial genome. In 2011, Deng et al. [30] classified a complete mitochondrial DNA dataset of 31 mammalian genome sequences from GenBank. The dataset was classified into 7 groups consisting of *Carnivore, Perissodactyla, Cetacea and Artiodactyla, Lagomorpha, Rodentia, Primates, and Erinaceomorpha*. In this subsection, we use the same datatset to test our method, based on edit distance and spectral distance. Using our method, based on edit distance (see Figure 6 and Tables 1 and 2), we correctly group 31 mammalian genome sequences into their corresponding 7 groups. Both Clustal Omega and the Fourier transform method in [24] have a misplacement. It took Clustal Omega 10 minutes and 9 seconds for the classification, and the Fourier transform method in [24] 4 seconds for the classification. It took our method using edit distance 4.34 seconds to correctly classify 31 mammalian genome sequences. Our method, based on spectral distance have two misplacements (see Figure 7), i.e. both Rabbit and Pig were misplaced in different groups. It took our method using spectral distance 42.60 seconds for classification.

**Figure 6:**
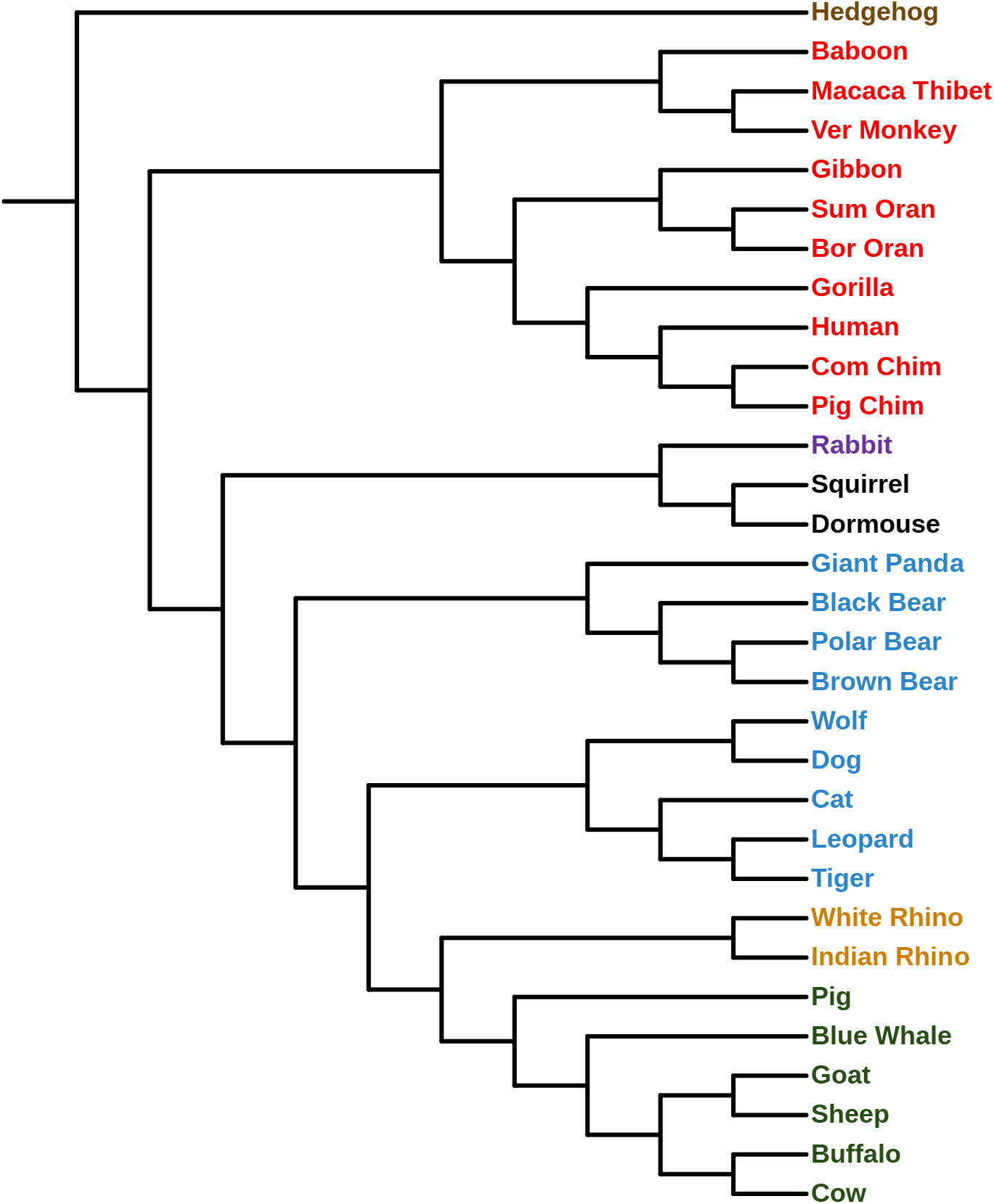
Phylogenetic tree of Mammals based on edit distance.

**Figure 7:**
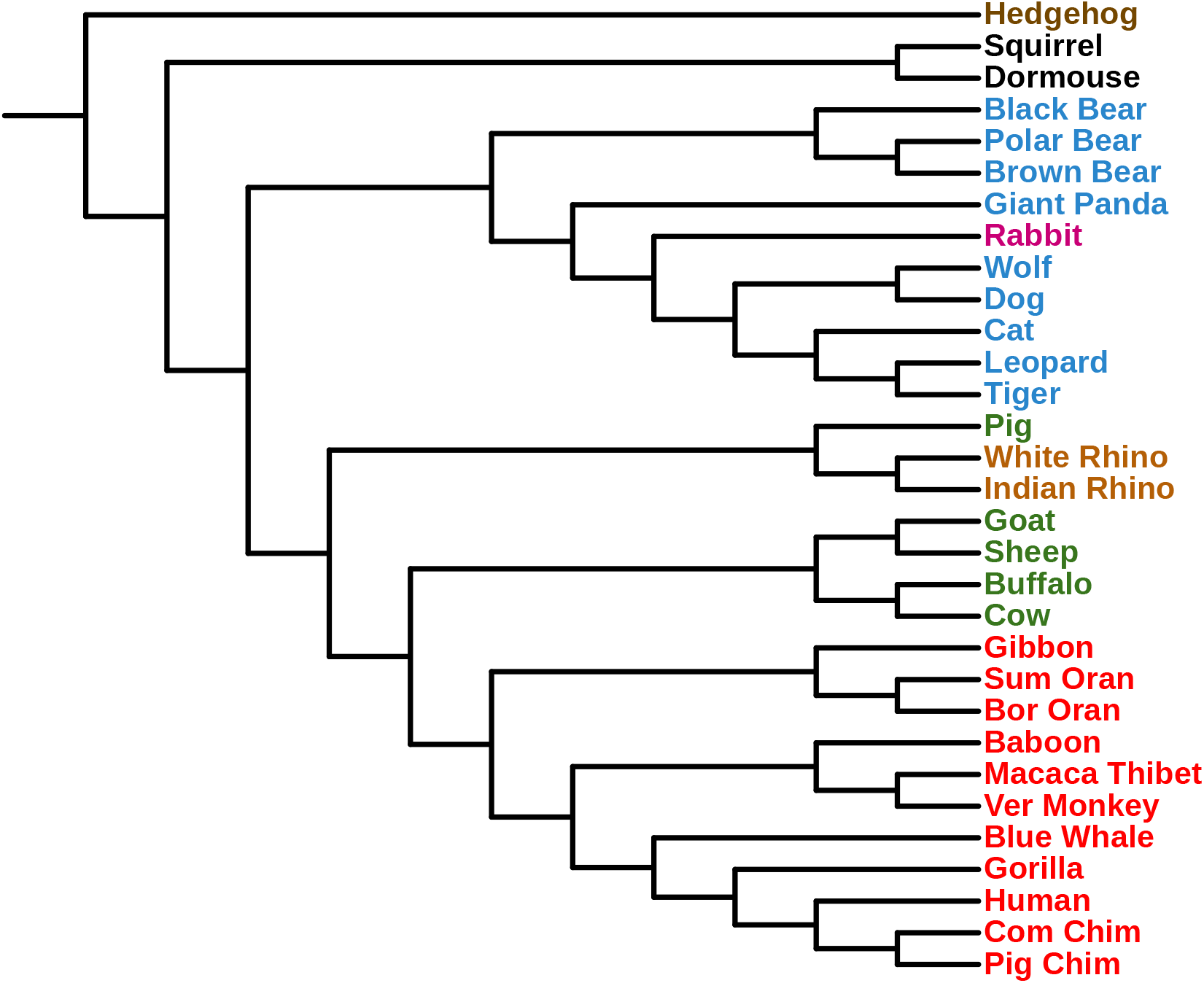
Phylogenetic tree of Mammals based on spectral distance.

### 4.4 Human rhinovirus (HRV)

Human rhinovirus (HRV) is the most common viral infectious agent in humans, and is the main cause of the common cold (see [27]). Using multiple sequence alignment, Palmenberg et al. [27] correctly classified the complete HRV genomes into three genetically distinct groups within the genus *Enterovirus* (HEV) and the family *Picornaviridae*. The dataset used in [27] consists of three groups HRV-A, HRV-B, HRV-C including 113 genomes, and three outgroup sequences HEV-C. Our method, using edit distance (see Tables 1 and 2 and Figure 8) for the phylogenetic tree of HRV genomes correctly classifies the complete HRV genomes into the corresponding genetically distinct groups in about 8.15 seconds. Our method, using edit distance, is comparable to the Fourier transform method in [24], which performs the computation in 7 seconds. Our method, based on spectral distance, incorrectly classifies one HRV genome (see the column HRV in Table 2). The phylogenetic tree of HRV genomes, based on spectral distance is illustrated in Figure 9.

**Figure 8:**
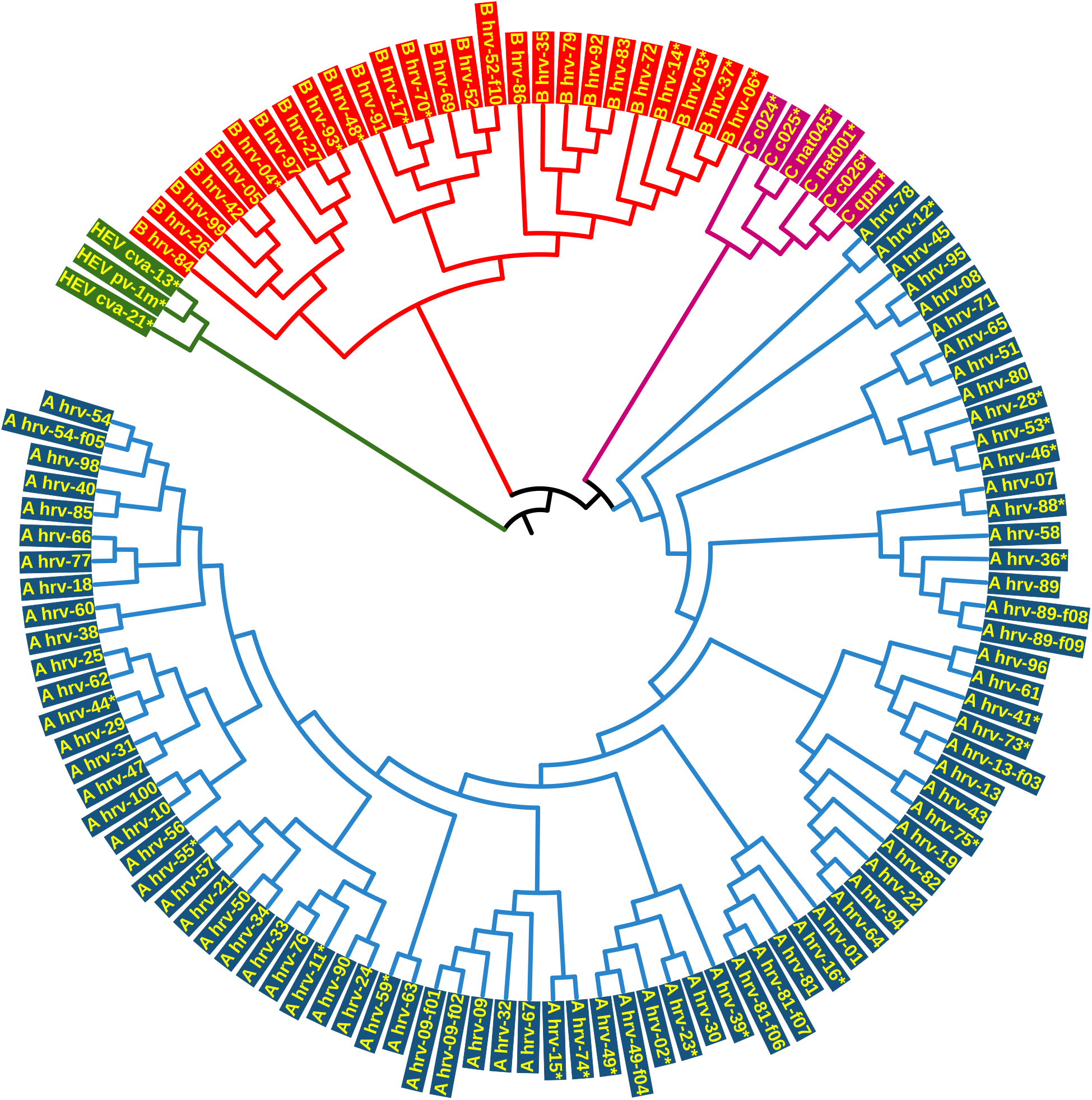
Phylogenetic tree of HRV based on edit distance.

**Figure 9:**
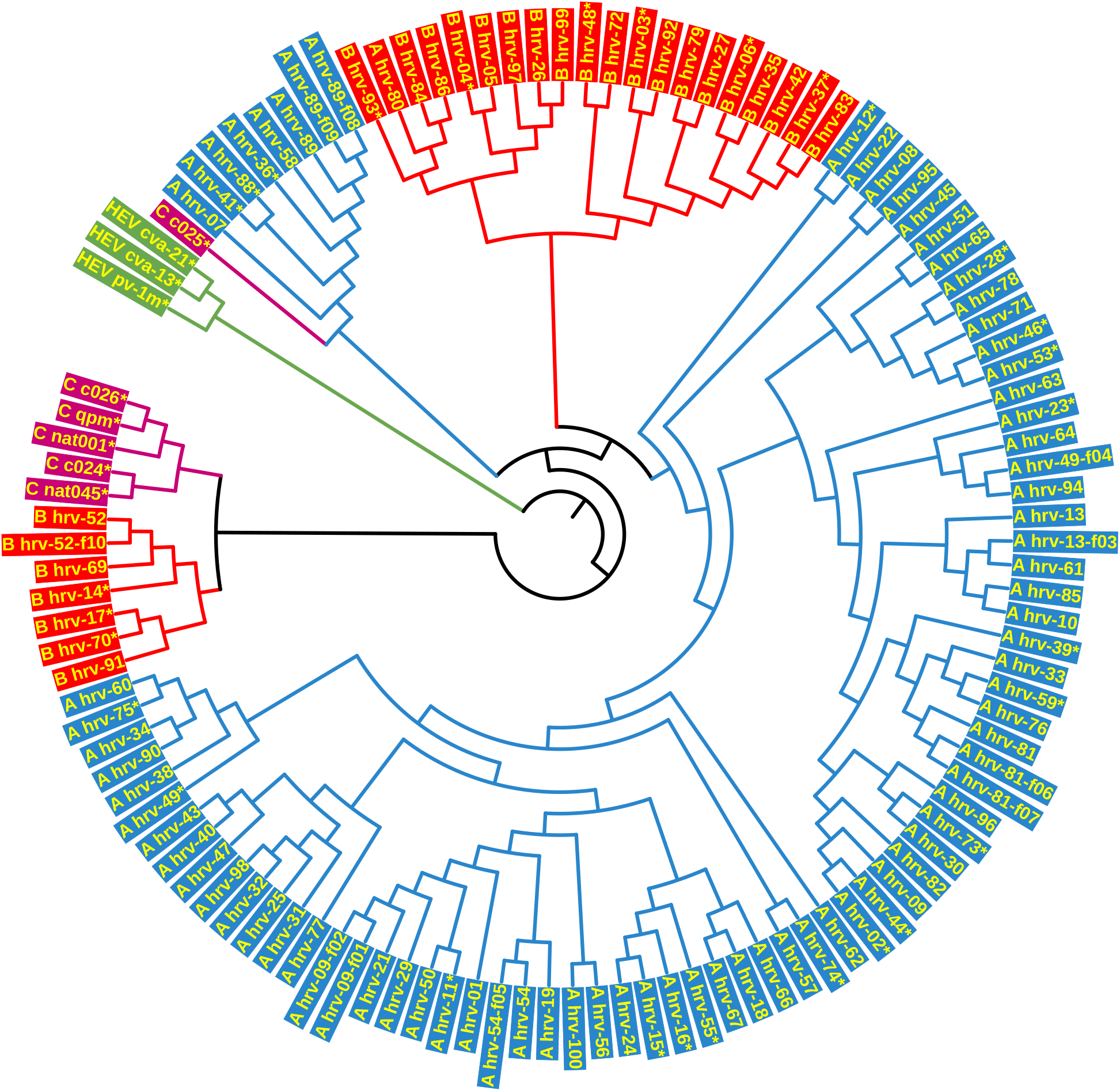
Phylogenetic tree of HRV based on spectral distance.

## 5 Appendix

**Table 3:** Information from GenBank of the 38 Influenza A viruses

| Accession Number | Description | Length |
| --- | --- | --- |
| HM370969 | A/turkey/Ontario/FAV110-4/2009(H1N1) | 1419 |
| CY138562 | A/mallard/Nova Scotia/00088/2010(H1N1) | 1422 |
| CY149630 | A/thick-billed murre/Canada/1871/2011(H1N1) | 1433 |
| KC608160 | A/duck/Guangxi/030D/2009(H1N1) | 1398 |
| AM157358 | A/mallard/France/691/2002(H1N1) | 1413 |
| AB470663 | A/duck/Hokkaido/w73/2007(H1N1) | 1422 |
| AB546159 | A/pintail/Miyagi/1472/2008(H1N1) | 1410 |
| HQ897966 | A/mallard/Korea/KNU YP09/2009(H1N1) | 1410 |
| EU026046 | A/mallard/Maryland/352/2002(H1N1) | 1433 |
| FJ357114 | A/mallard/Maryland/26/2003(H1N1) | 1433 |
| GQ411894 | A/dunlin/Alaska/44421-660/2008(H1N1) | 1413 |
| CY140047 | A/mallard/Minnesota/Sg-00620/2008(H1N1) | 1433 |
| KM244078 | A/turkey/Virginia/4135/2014(H1N1) | 1410 |
| HQ185381 | A/chicken/Eastern China/XH222/2008(H5N1) | 1350 |
| HQ185383 | A/duck/Eastern China/JS017/2009(H5N1) | 1350 |
| EU635875 | A/chicken/Yunnan/chuxiong01/2005(H5N1) | 1350 |
| FM177121 | A/chicken/Germany/R3234/2007(H5N1) | 1370 |
| AM914017 | A/domestic duck/Germany/R1772/2007(H5N1) | 1350 |
| KF572435 | A/wild bird/Hong Kong/07035-1/2011(H5N1) | 1350 |
| AF509102 | A/Chicken/Hong Kong/822.1/01 (H5N1) | 1366 |
| AB684161 | A/chicken/Miyazaki/10/2011(H5N1) | 1350 |
| EF541464 | A/chicken/Korea/es/2003(H5N1) | 1350 |
| JF699677 | A/mandarin duck/Korea/K10-483/2010(H5N1) | 1350 |
| GU186511 | A/turkey/VA/505477-18/2007(H5N1) | 1370 |
| EU500854 | A/American black duck/NB/2538/2007(H7N3) | 1453 |
| CY129336 | A/American black duck/New Brunswick/02490/2007(H7N3) | 1428 |
| CY076231 | A/American green-winged teal/California/44242-906/2007(H7N3) | 1420 |
| CY039321 | A/avian/Delaware Bay/226/2006(H7N3) | 1434 |
| AY646080 | A/chicken/British Columbia/GSC_human_B/04(H7N3) | 1453 |
| KF259734 | A/chicken/Rizhao/713/2013(H7N9) | 1398 |
| KF938945 | A/chicken/Jiangsu/1021/2013(H7N9) | 1404 |
| KF259688 | A/duck/Jiangxi/3096/2009(H7N9) | 1413 |
| KC609801 | A/wild duck/Korea/SH19-47/2010(H7N9) | 1426 |
| CY014788 | A/turkey/Minnesota/1/1988(H7N9) | 1460 |
| CY186004 | A/mallard/Minnesota/AI09-3770/2009(H7N9) | 1422 |
| DQ017487 | A/mallard/Postdam/178-4/1983(H2N2) | 1467 |
| CY005540 | A/duck/Hong Kong/319/1978(H2N2) | 1467 |
| JX081142 | A/emperor goose/Alaska/44297-260/2007(H2N2) | 1457 |

**Table 4:**
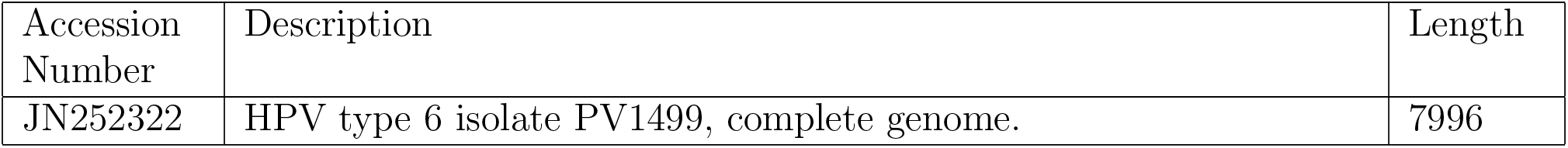
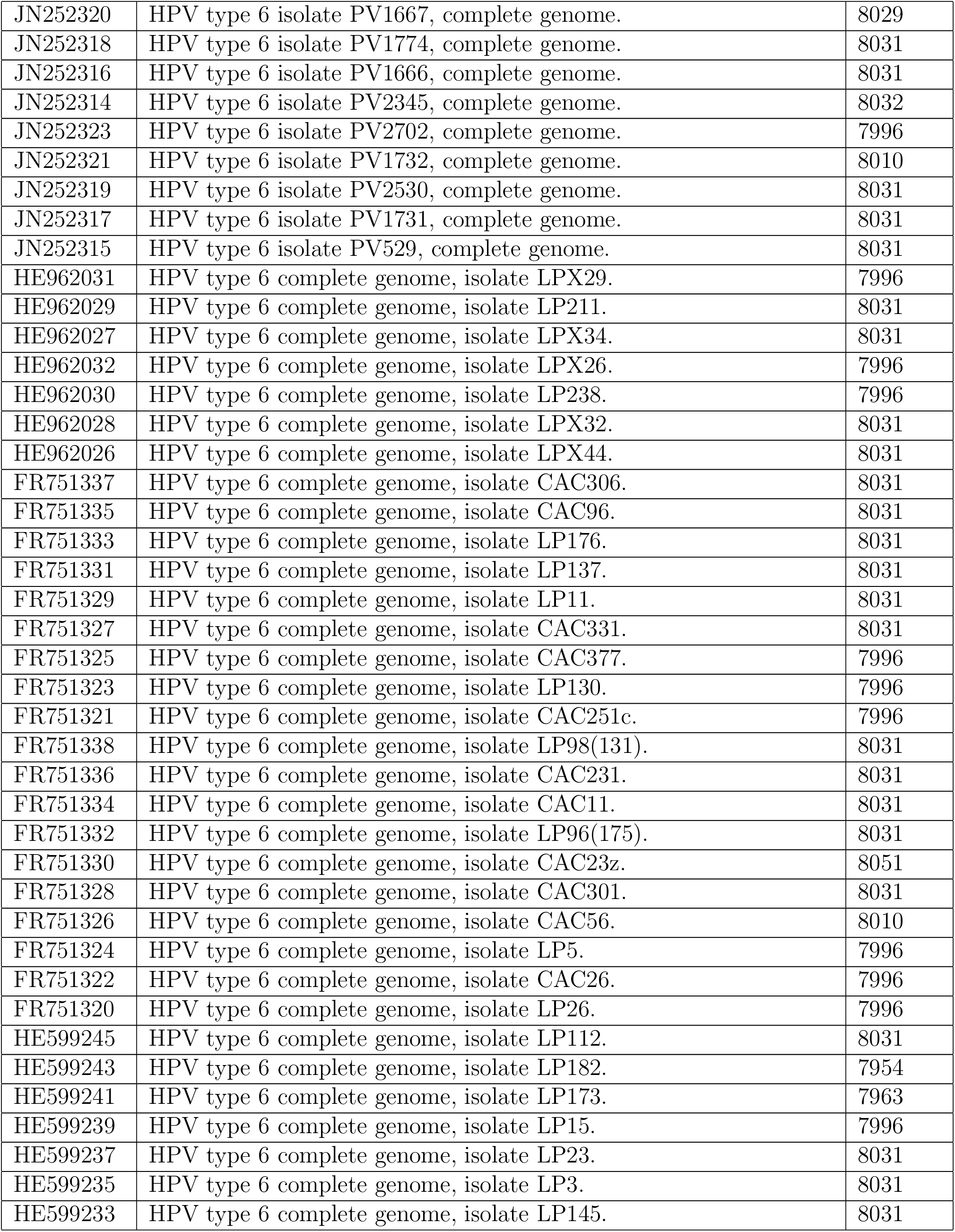
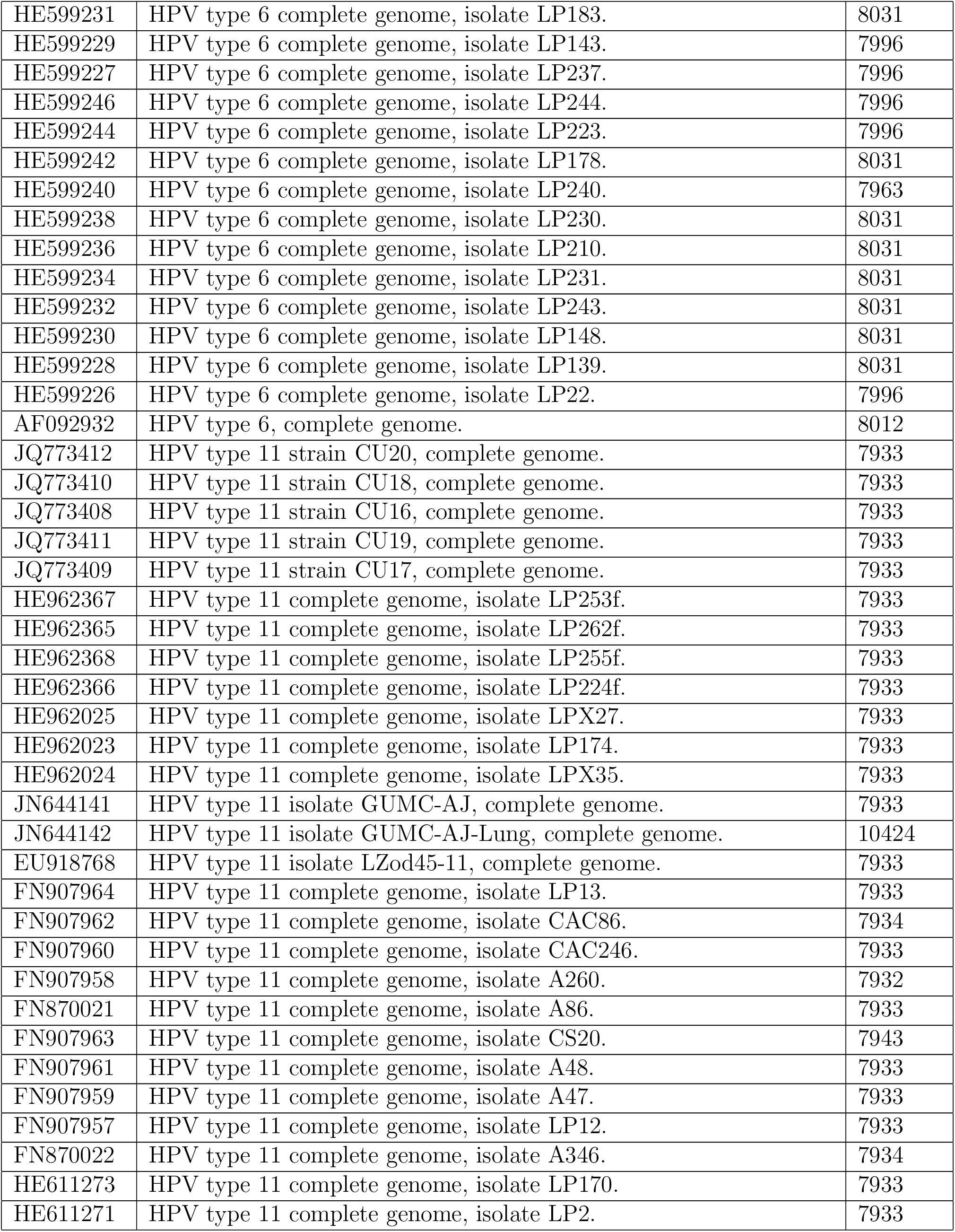
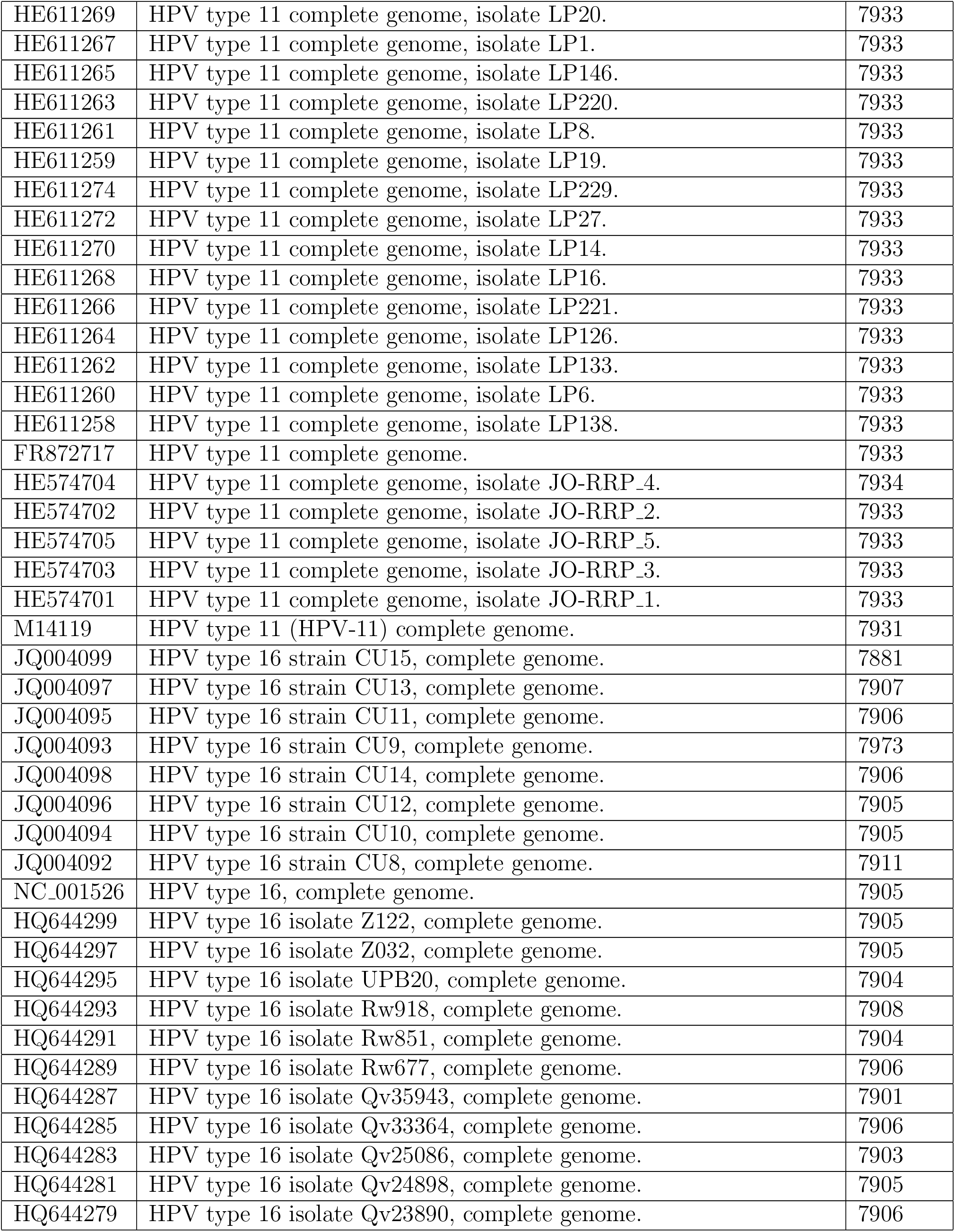
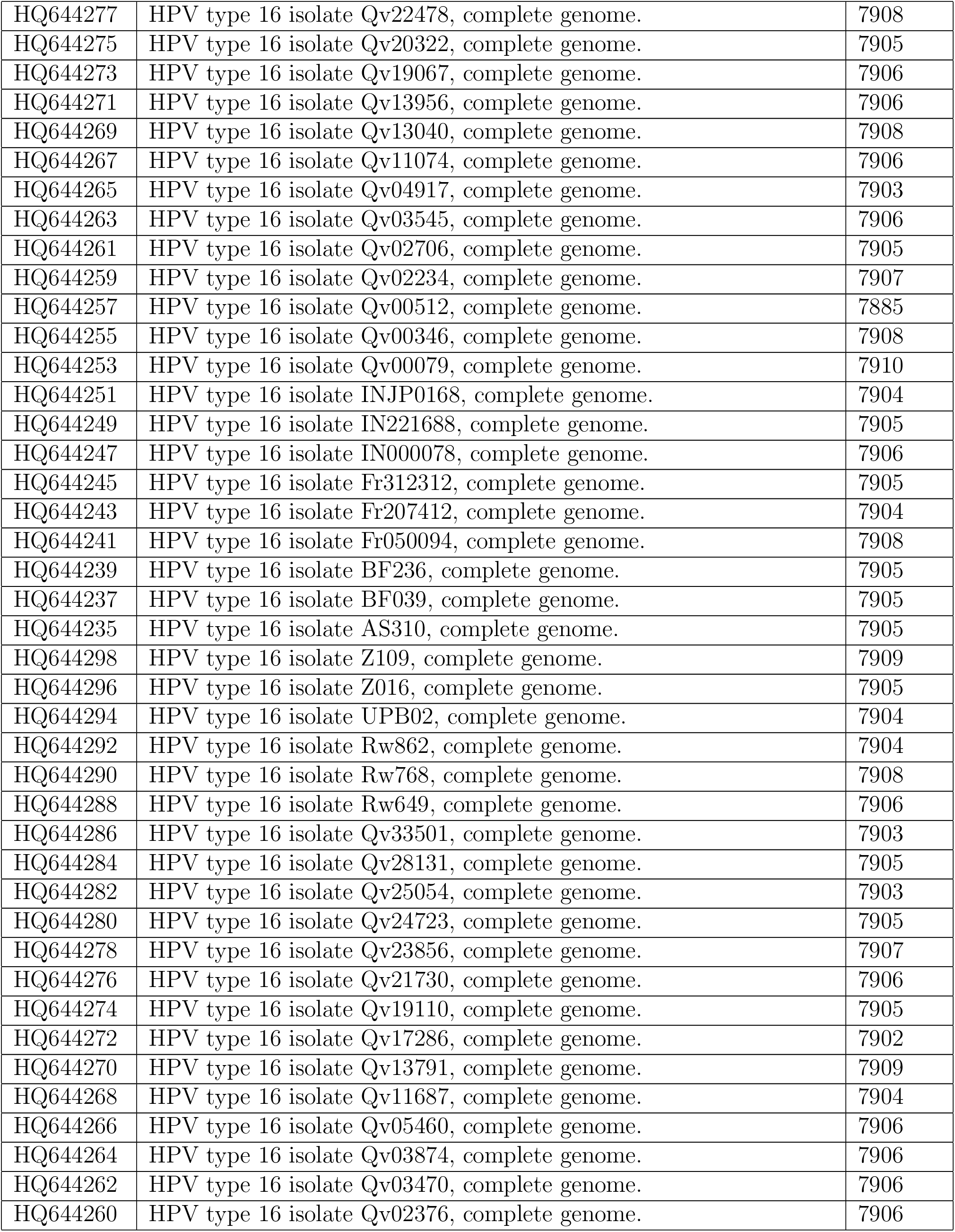
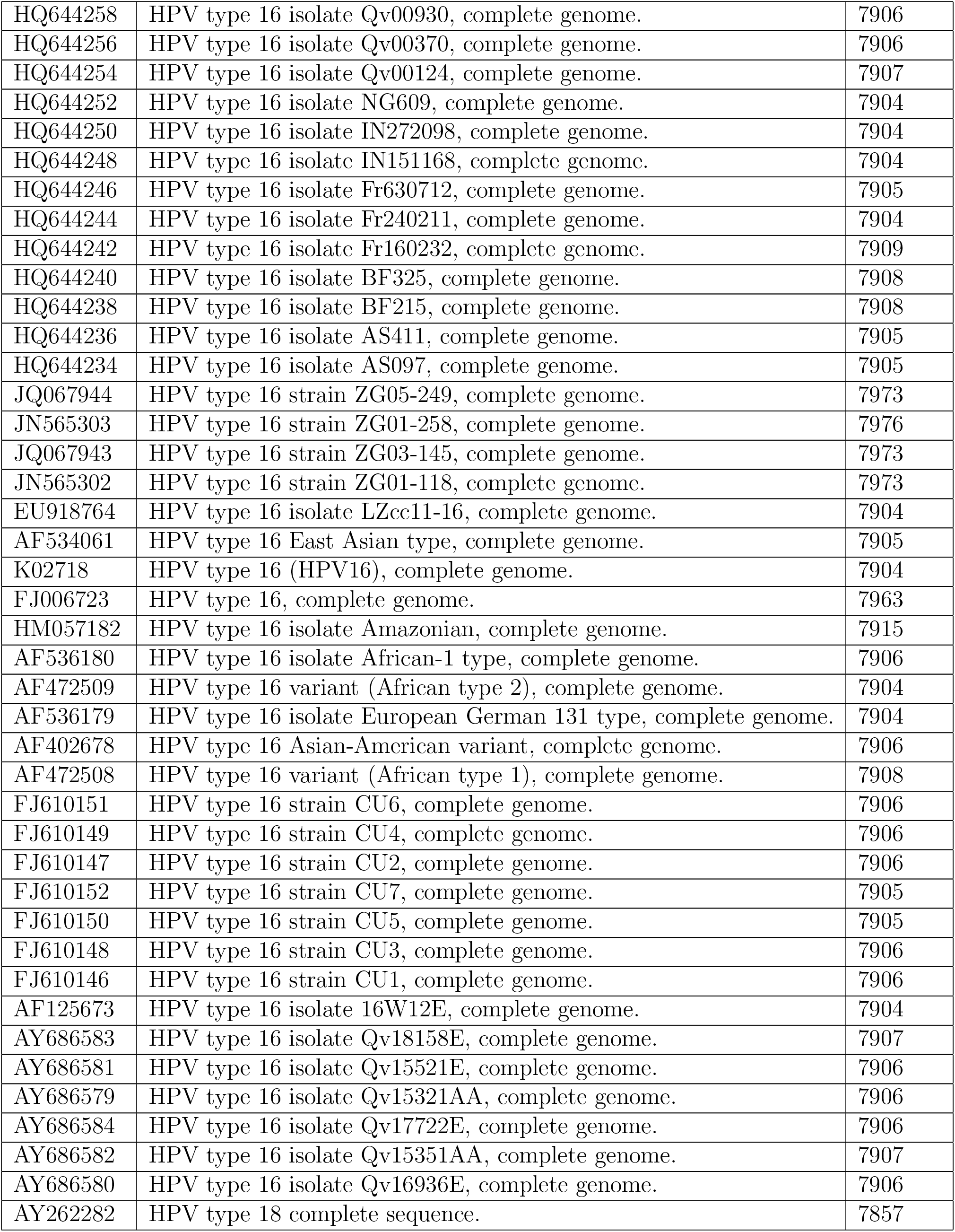
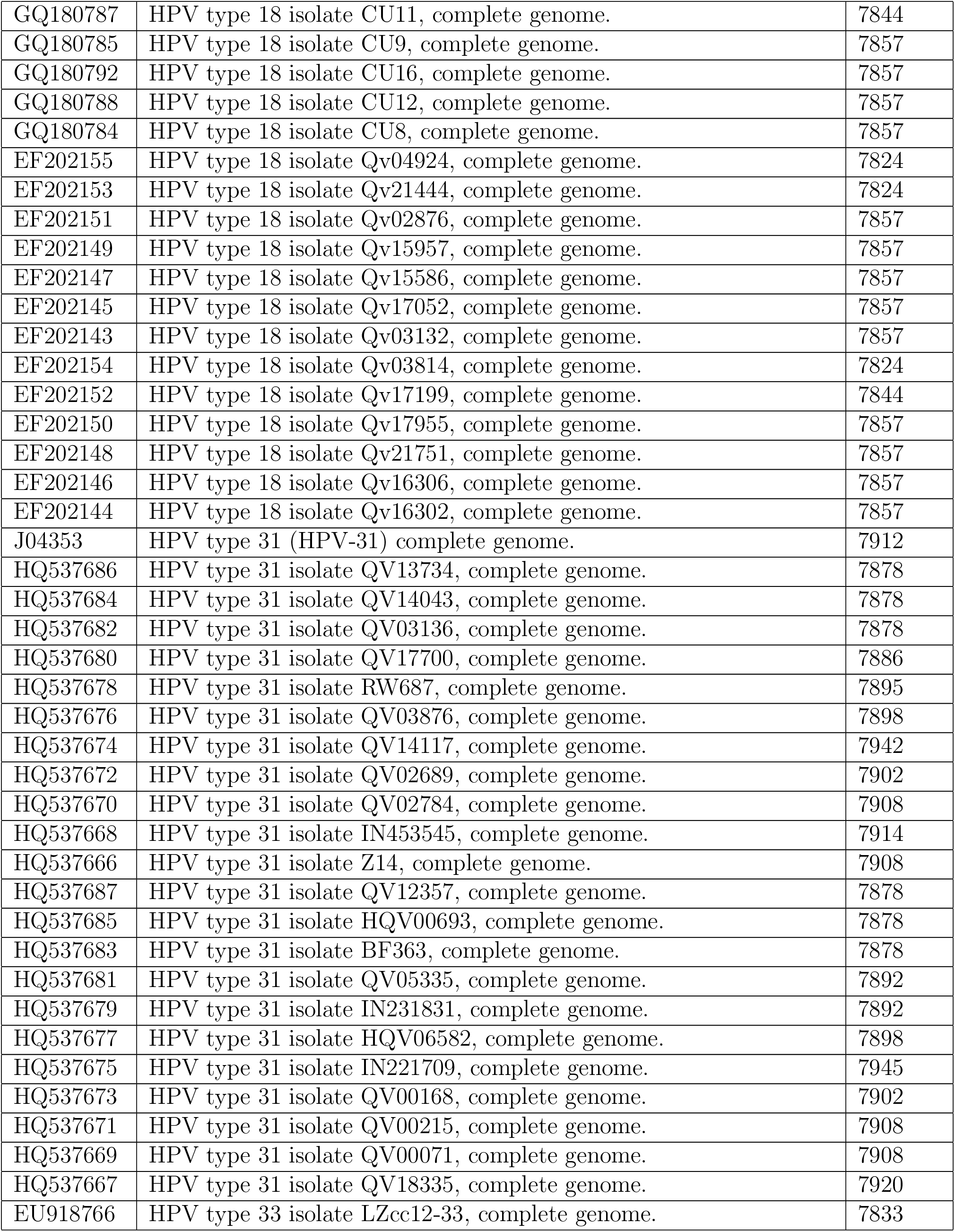
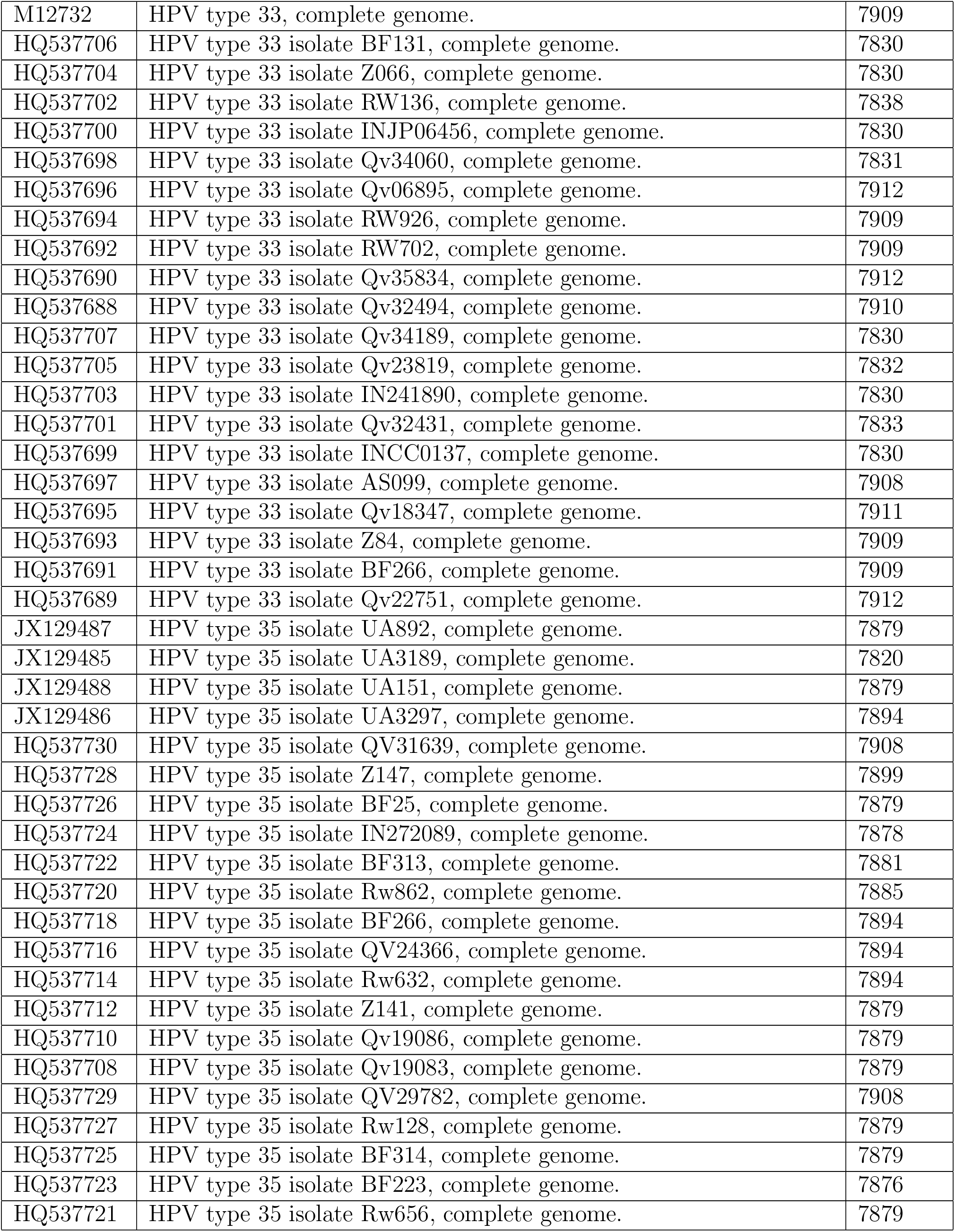
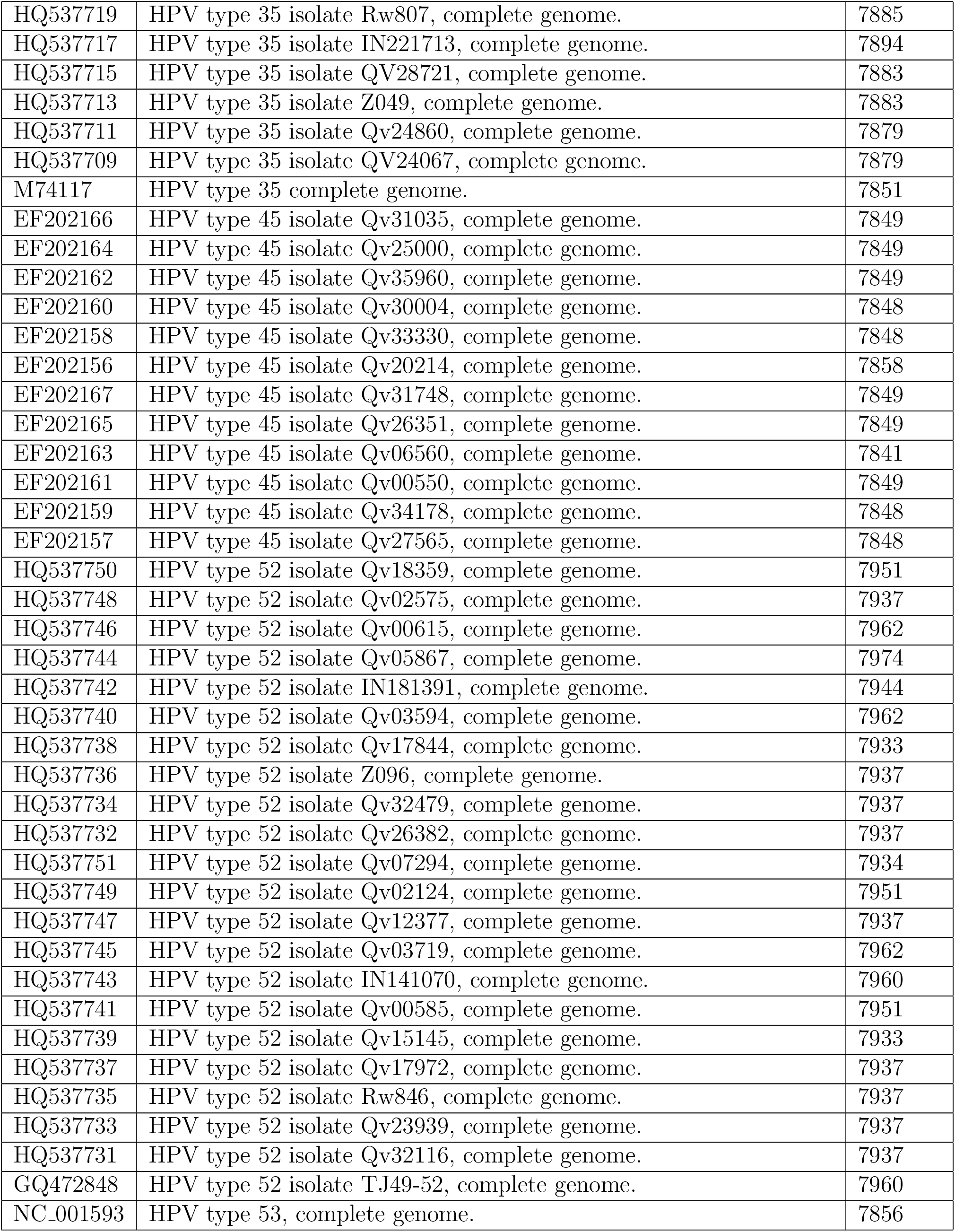
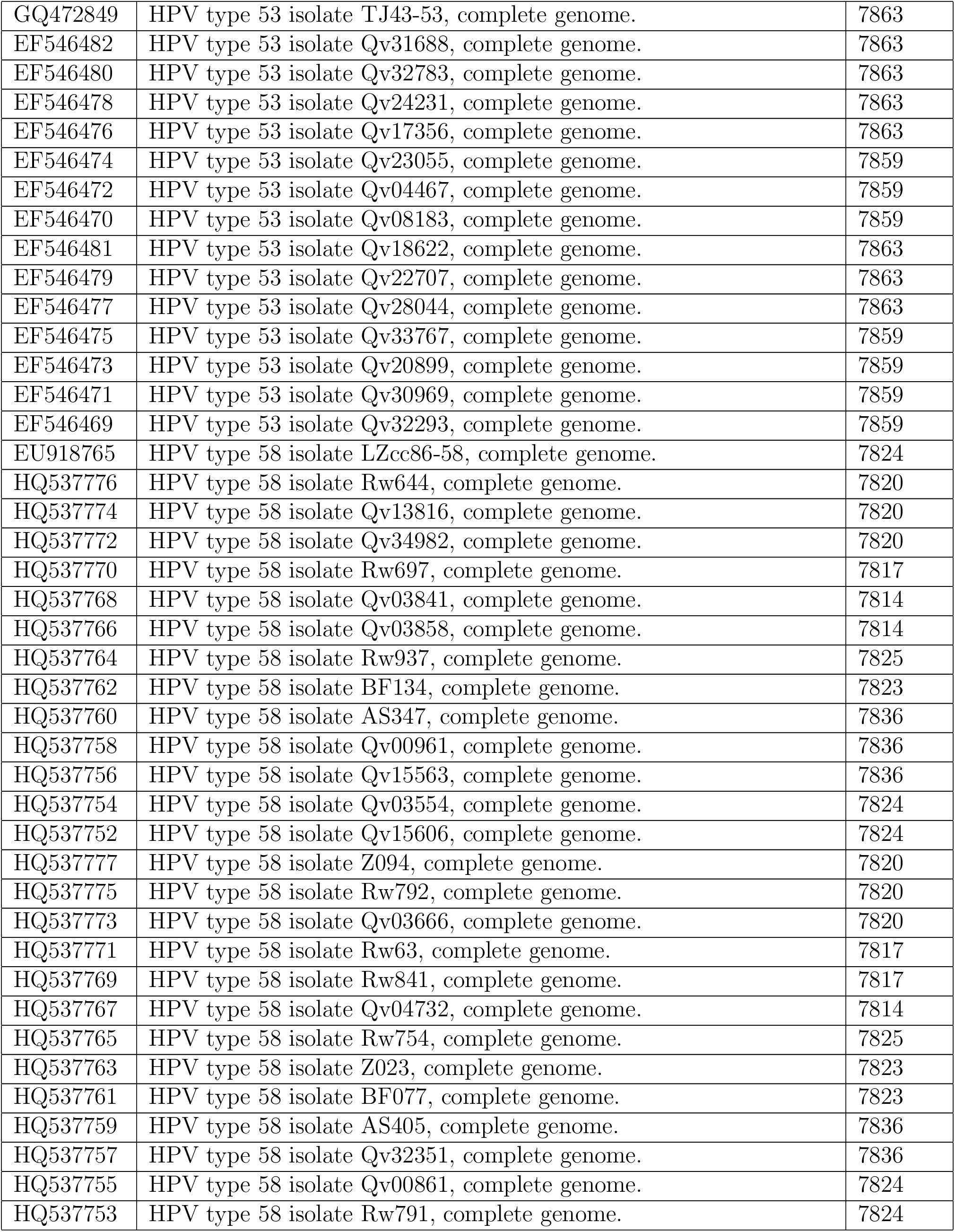
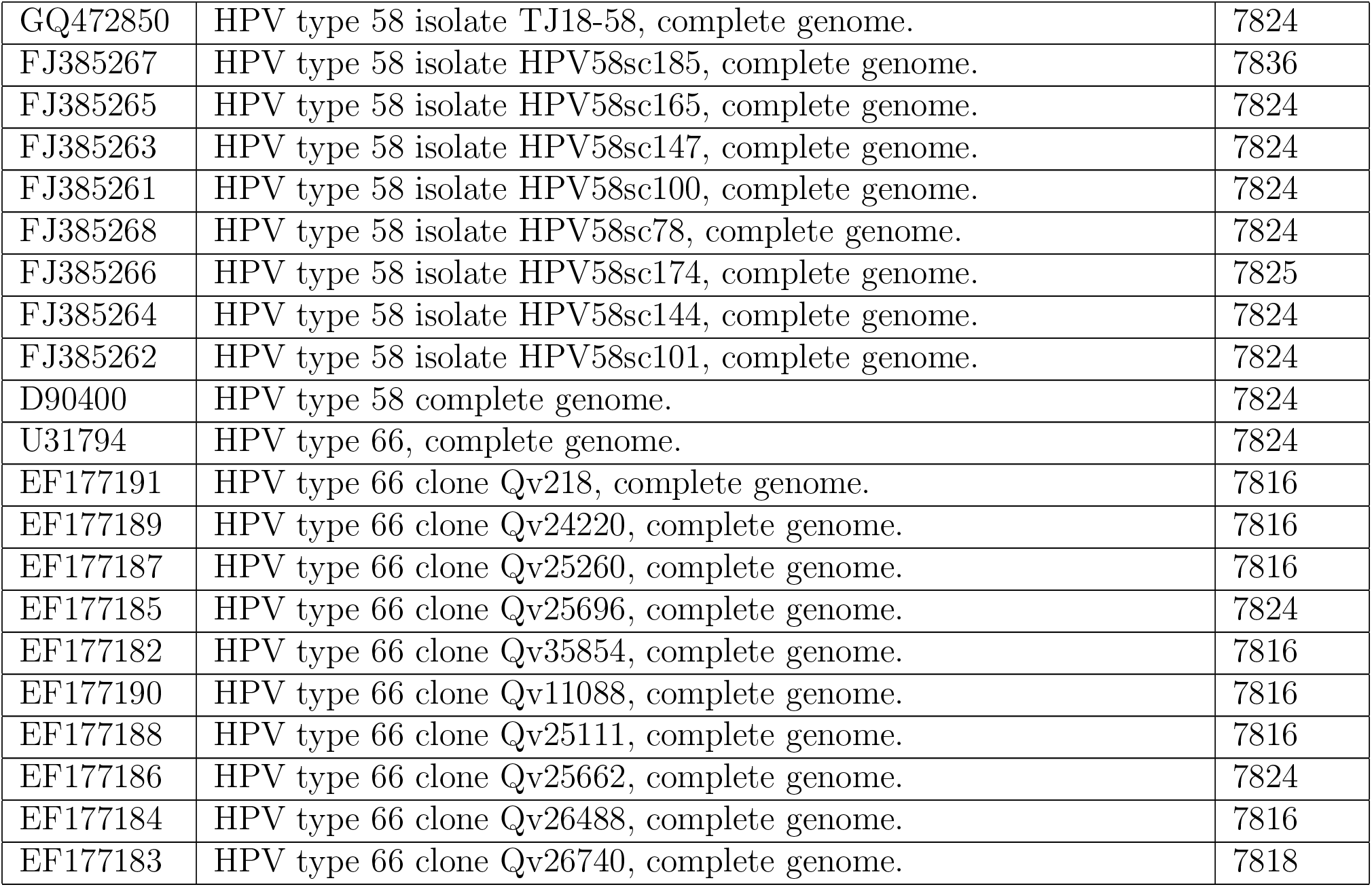
Information from GenBank of 400 HPV data set.

**Table 5:** Information from GenBank of 113 human rhinovirus genomes and three outgroup genomes

| Accession Number | Abbreviation | Length |
| --- | --- | --- |
| AF499637 | HEV_cva-13* | 7458 |
| AF546702 | HEV_cva-21* | 7406 |
| AY751783 | A_hrv-39* | 7137 |
| DQ473485 | B_hrv-03* | 7208 |
| DQ473486 | B_hrv-06* | 7216 |
| DQ473488 | B_hrv-48* | 7214 |
| DQ473489 | B_hrv-70* | 7223 |
| DQ473490 | B_hrv-04* | 7212 |
| DQ473491 | A_hrv-41* | 7145 |
| DQ473492 | A_hrv-73* | 7140 |
| DQ473493 | A_hrv-15* | 7134 |
| DQ473494 | A_hrv-74* | 7120 |
| DQ473496 | A_hrv-49* | 7106 |
| DQ473497 | A_hrv-23* | 7025 |
| DQ473499 | A_hrv-44* | 7123 |
| DQ473500 | A_hrv-59* | 7135 |
| DQ473504 | A_hrv-88* | 7143 |
| DQ473505 | A_hrv-36* | 7141 |
| DQ473506 | A_hrv-46* | 7149 |
| DQ473507 | A_hrv-53* | 7143 |
| DQ473508 | A_hrv-28* | 7148 |
| DQ473510 | A_hrv-75* | 7137 |
| DQ473511 | A_hrv-55* | 7036 |
| EF077279 | C_nat001* | 6944 |
| EF077280 | C_nat045* | 7015 |
| EF173414 | A_hrv-11* | 7125 |
| EF173415 | A_hrv-12* | 7124 |
| EF173420 | B_hrv-17* | 7219 |
| EF173423 | B_hrv-37* | 7216 |
| EF173425 | B_hrv-93* | 7215 |
| EF186077 | C_qpm* | 7134 |
| EF582385 | C_c024* | 7099 |
| EF582386 | C_c025* | 7114 |
| EF582387 | C_c026* | 7086 |
| FJ445111 | A_hrv-01 | 7137 |
| FJ445112 | B_hrv-05 | 7212 |
| FJ445113 | A_hrv-08 | 7108 |
| FJ445114 | A_hrv-09-f01 | 7134 |
| FJ445115 | A_hrv-09-f02 | 7133 |
| FJ445116 | A_hrv-13 | 7140 |
| FJ445117 | A_hrv-13-f03 | 7143 |
| FJ445118 | A_hrv-18 | 7119 |
| FJ445119 | A_hrv-19 | 7135 |
| FJ445121 | A_hrv-21 | 7134 |
| FJ445122 | A_hrv-22 | 7129 |
| FJ445123 | A_hrv-25 | 7126 |
| FJ445124 | B_hrv-26 | 7211 |
| FJ445125 | A_hrv-29 | 7123 |
| FJ445126 | A_hrv-31 | 7131 |
| FJ445127 | A_hrv-32 | 7133 |
| FJ445128 | A_hrv-33 | 7133 |
| FJ445129 | A_hrv-40 | 7138 |
| FJ445130 | B_hrv-42 | 7223 |
| FJ445131 | A_hrv-43 | 7129 |
| FJ445132 | A_hrv-45 | 7114 |
| FJ445133 | A_hrv-47 | 7132 |
| FJ445134 | A_hrv-49-f04 | 7109 |
| FJ445135 | A_hrv-50 | 7118 |
| FJ445136 | A_hrv-51 | 7152 |
| FJ445137 | B_hrv-52-f10 | 7216 |
| FJ445138 | A_hrv-54 | 7134 |
| FJ445139 | A_hrv-54-f05 | 7133 |
| FJ445140 | A_hrv-56 | 7136 |
| FJ445141 | A_hrv-57 | 7134 |
| FJ445142 | A_hrv-58 | 7140 |
| FJ445143 | A_hrv-60 | 7139 |
| FJ445144 | A_hrv-61 | 7139 |
| FJ445145 | A_hrv-62 | 7131 |
| FJ445146 | A_hrv-63 | 7141 |
| FJ445147 | A_hrv-65 | 7162 |
| FJ445148 | A_hrv-66 | 7139 |
| FJ445149 | A_hrv-67 | 7135 |
| FJ445151 | B_hrv-69 | 7211 |
| FJ445152 | A_hrv-71 | 7161 |
| FJ445153 | B_hrv-72 | 7216 |
| FJ445154 | A_hrv-77 | 7136 |
| FJ445155 | B_hrv-79 | 7224 |
| FJ445156 | A_hrv-80 | 7138 |
| FJ445157 | A_hrv-81 | 7116 |
| FJ445158 | A_hrv-81-f06 | 7116 |
| FJ445159 | A_hrv-81-f07 | 7116 |
| FJ445160 | A_hrv-82 | 7123 |
| FJ445161 | B_hrv-83 | 7230 |
| FJ445162 | B_hrv-84 | 7201 |
| FJ445163 | A_hrv-85 | 7140 |
| FJ445164 | B_hrv-86 | 7213 |
| FJ445165 | A_hrv-89-f09 | 7150 |
| FJ445166 | A_hrv-89-f08 | 7152 |
| FJ445167 | A_hrv-90 | 7124 |
| FJ445168 | B_hrv-91 | 7221 |
| FJ445169 | B_hrv-92 | 7233 |
| FJ445170 | A_hrv-95 | 7110 |
| FJ445171 | A_hrv-96 | 7134 |
| FJ445172 | B_hrv-97 | 7207 |
| FJ445173 | A_hrv-98 | 7133 |
| FJ445174 | B_hrv-99 | 7208 |
| FJ445175 | A_hrv-100 | 7140 |
| FJ445176 | A_hrv-07 | 7146 |
| FJ445177 | A_hrv-09 | 7132 |
| FJ445178 | A_hrv-10 | 7137 |
| FJ445179 | A_hrv-30 | 7093 |
| FJ445180 | A_hrv-38 | 7136 |
| FJ445181 | A_hrv-64 | 7129 |
| FJ445182 | A_hrv-76 | 7128 |
| FJ445183 | A_hrv-78 | 7145 |
| FJ445184 | A_hrv-89 | 7152 |
| FJ445185 | A_hrv-94 | 7132 |
| FJ445186 | B_hrv-27 | 7217 |
| FJ445187 | B_hrv-35 | 7224 |
| FJ445188 | B_hrv-52 | 7216 |
| FJ445189 | A_hrv-34 | 7119 |
| FJ445190 | A_hrv-24 | 7132 |
| L05355 | B_hrv-14* | 7212 |
| L24917 | A_hrv-16* | 7124 |
| V01149 | HEV_pv-1m* | 7440 |
| X02316 | A_hrv-02* | 7102 |

**Table 6:** Information from GeneBank of 31 mammalian mitochondrial genomes

| GeneBank ID | Name | Abbreviation | Length |
| --- | --- | --- | --- |
| V00662 | Human | Human | 16569 |
| D38113 | Common chimpanzee | Com Chim | 16554 |
| D38116 | Pigmy chimpanzee | Pig Chim | 16563 |
| D38114 | Gorilla | Gorilla | 16472 |
| N_002764 | Macaca thibetana | Macaca Thibet | 17447 |
| D38115 | Bornean Orangutan | Bor Oran | 16389 |
| NC_002083 | Sumatran Orangutan | Sum Oran | 16499 |
| X99256 | Gibbon | Gibbon | 16521 |
| AY863426 | Vervet Monkey | Ver Monkey | 16586 |
| Y18001 | Baboon | Baboon | 16389 |
| U20753 | Cat | Cat | 16364 |
| U96639 | Dog | Dog | 17009 |
| V00654 | Cow | Cow | 16640 |
| AF010406 | Sheep | Sheep | 26680 |
| AF533441 | Goat | Goat | 16616 |
| AY488491 | Buffalo | Buffalo | 16638 |
| AJ002189 | Pig | Pig | 16727 |
| EU442884 | Wolf | Wolf | 16355 |
| DQ402478 | Black bear | Black bear | 16832 |
| AF303111 | Polar bear | Polar bear | 17020 |
| AF303110 | Brown bear | Brown bear | 16868 |
| EF212882 | Giant panda | Giant panda | 17017 |
| X97336 | Indian rhinoceros | Indian Rhino | 16964 |
| Y07726 | White rhinoceros | White Rhino | 16829 |
| AJ001588 | Rabbit | Rabbit | 16805 |
| EF551003 | Tiger | Tiger | 16774 |
| EF551002 | Leopard | Leopard | 16990 |
| AJ001562 | Dormouse | Dormouse | 16507 |
| X72204 | Blue Whale | Blue Whale | 16402 |
| AJ238588 | Squirrel | Squirrel | 16602 |
| X88898 | Hedgehog | Hedgehog | 17245 |

## Footnotes

* See https://www.ncbi.nlm.nih.gov/genbank/

* https://github.com/peterewills/netcomp

† See https://github.com/peterewills/netcomp

‡ See https://www.ncbi.nlm.nih.gov/genbank/

† a part of H1N1 viruses are incorrectly grouped with H5N1 subtype

## References

[1] X. Jin, R. Nie, D. Zhou, S. Yao, Y. Chen, J. Yu, and Q. Wang, “A novel dna sequence similarity calculation based on simplified pulse-coupled neural network and huffman coding,” Physica A: Statistical Mechanics and its Applications, vol. 461, pp. 325–338, 2016.

[2] J. F. Yu, J. H. Wang, and X. Sun, “Analysis of similarities/dissimilarities of dna sequences based on a novel graphical representation,” MATCH Commun. Math. Comput. Chem, vol. 63, no. 2, pp. 493–512, 2010.

[3] S. Wang, F. Tian, Y. Qiu, and X. Liu, “Bilateral similarity function: A novel and universal method for similarity analysis of biological sequences,” Journal of theoretical biology, vol. 265, no. 2, pp. 194–201, 2010.

[4] N. Jafarzadeh and A. Iranmanesh, “C-curve: a novel 3d graphical representation of dna sequence based on codons,” Mathematical Biosciences, vol. 241, no. 2, pp. 217–224, 2013.

[5] B. Liao, Y. Zhang, K. Ding, and T. M. Wang, “Analysis of similarity/dissimilarity of dna sequences based on a condensed curve representation,” Journal of Molecular Structure: THEOCHEM, vol. 717, no. 1-3, pp. 199–203, 2005.

[6] E. Hamori and J. Ruskin, “H curves, a novel method of representation of nucleotide series especially suited for long dna sequences.,” Journal of Biological Chemistry, vol. 258, no. 2, pp. 1318–1327, 1983.

[7] B. Liao, M. Tan, and K. Ding, “A 4d representation of dna sequences and its application,” Chemical Physics Letters, vol. 402, no. 4-6, pp. 380–383, 2005.

[8] J. Wang and Y. Zhang, “Characterization and similarity analysis of dna sequences grounded on a 2-d graphical representation,” Chemical physics letters, vol. 423, no. 1-3, pp. 50–53, 2006.

[9] X. Q. Liu, Q. Dai, Z. Xiu, and T. Wang, “Pnn-curve: A new 2d graphical representation of dna sequences and its application,” Journal of Theoretical Biology, vol. 243, no. 4, pp. 555–561, 2006.

[10] X. Q. Qi, J. Wen, and Z. H. Qi, “New 3d graphical representation of dna sequence based on dual nucleotides,” Journal of Theoretical Biology, vol. 249, no. 4, pp. 681–690, 2007.

[11] C. Li, X. Yu, and N. Helal, “Similarity analysis of dna sequences based on codon usage,” Chemical Physics Letters, vol. 459, no. 1-6, pp. 172–174, 2008.

[12] N. Jafarzadeh and A. Iranmanesh, “A novel graphical and numerical representation for analyzing dna sequences based on codons,” Match-Communications in Mathematical and Computer Chemistry, vol. 68, no. 2, p. 611, 2012.

[13] F. Kabli, H. R. Mohamed, and A. Abdelmalek, “Similarity analysis of dna sequences based on the chemical properties of nucleotide bases: frequency and position of group mutations,” Comput. Sci. Inf. Technol., vol. 6, no. 1, pp. 1–10, 2016.

[14] Q. Dai, X. Liu, and T. Wang, “A novel 2d graphical representation of dna sequences and its application,” Journal of Molecular Graphics and Modelling, vol. 25, no. 3, pp. 340–344, 2006.

[15] Y. H. Yao, X. Y. Nan, and T. M. Wang, “A new 2d graphical representation— classification curve and the analysis of similarity/dissimilarity of dna sequences,” Journal of Molecular Structure: THEOCHEM, vol. 764, no. 1-3, pp. 101–108, 2006.

[16] B. Liao, Q. Xiang, L. Cai, and Z. Cao, “A new graphical coding of dna sequence and its similarity calculation,” Physica A: Statistical Mechanics and its Applications, vol. 392, no. 19, pp. 4663–4667, 2013.

[17] P. A. He and J. Wang, “Characteristic sequences for dna primary sequence,” Journal of Chemical Information & Modeling, vol. 42, no. 5, pp. 1080–1085, 2002.

[18] W. Hou, Q. Pan, and M. He, “A novel representation of dna sequence based on cmi coding,” Physica A, vol. 409, pp. 87–96, 2014.

[19] R. Zhang and C. Zhang, “Fast, scalable generation of high-quality protein multiple sequence alignments using clustal omega,” J Biomol Struct Dyn, vol. 11, no. 4, pp. 767–782, 1994.

[20] R. Zhang and C. Zhang, “A brief review: The z-curve theory and its application in genome analysis,” Curr Genomics, vol. 15, no. 2, pp. 78–94, 2014.

[21] P. Wills and F. Meyer, “Metrics for graph comparison: A practitioner’s guide,” PLOS ONE, vol. 15, pp. 1–54, 02 2020.

[22] J. A. Dieudonné, Foundations of modern analysis. Pure and applied mathematics (Academic Press) ; 10, New York: Academic Press, 1960.

[23] F. Sievers, A. Wilm, D. Dineen, T. Gibson, K. Karplus, W. Li, R. Lopez, H. McWilliam, M. Remmert, and e. a. J. Söding, “Fast, scalable generation of high-quality protein multiple sequence alignments using clustal omega,” Mol. Syst. Biol., vol. 7, p. 539, 2011.

[24] T. Hoang, C. Yin, and S. S.-T. Yau, “Numerical encoding of dna sequences by chaos game representation with application in similarity comparison.,” Genomics, vol. 108, no. 3-4, pp. 134–142, 2016.

[25] M. Arbyn, X. Castellsague, S. D. Sanjose, L. Bruni, M. Saraiya, F. Bray, and J. Ferlay, “Worldwide burden of cervical cancer in 2008,” Ann. Oncol., vol. 22, pp. 2675–2686, 2011.

[26] J. Smith, L. Lindsay, B. Hoots, J. Keys, S. Franceschi, R. Winer, and G. Clifford, “Human papillomavirus type distribution in invasive cervical cancer and high-grade cervical lesions: a meta-analysis update,” Int. J. Cancer, vol. 121, pp. 621–632, 2007.

[27] A. C. Palmenberg, D. Spiro, R. Kuzmickas, S. Wang, A. Djikengand, J. A. Ratheand, C. M. Fraser-Liggett, and L. S. b, “Sequencing and analyses of all known human rhinovirus genomes reveal structure and evolution,” Science (American Association for the Advancement of Science), vol. 324, no. 5923, pp. 55–59, 2009.

[28] R. Garten, C. Davis, C. Russell, B. Shu, S. Lindstrom, A. Balish, W. Sessions, E. S. X. Xu, and e. a. V. Deyde, “Antigenic and genetic characteristics of swine-origin 2009 a (h1n1) influenza viruses circulating in humans,” Science, vol. 325, pp. 197–201, 2009.

[29] P. Palese and J. Young, “Variation of influenza a, b, and c viruses,” Science, vol. 215, pp. 1468–1474, 1982.

[30] C. Yu, M. Deng, and S. S.-T. Yau, “Dna sequence comparison by a novel probabilistic method,” Information Sciences, vol. 181, no. 8, pp. 1484–1492, 2011.

[31] S. Kumar, G. Stecher, and K. Tamura, “MEGA7: Molecular Evolutionary Genetics Analysis Version 7.0 for Bigger Datasets,” Molecular Biology and Evolution, vol. 33, pp. 1870–1874, 03 2016.

[32] D. Alexander, “A review of avian influenza in different bird species,” Vet. Microbiol., vol. 74, pp. 3–13, 2000.

